# Zona incerta drives a strategy of persistence during nocturnal predatory hunting in mice

**DOI:** 10.64898/2026.11.13.766580

**Authors:** M.C. Shallow, A. Wright, A Weible, M Weible, M. Smear, M. Wehr

## Abstract

Prey capture in lab mice is a complex, ethologically relevant behavior that provides us with an opportunity to study how mice integrate internal and external cues to drive goal-directed behaviors. Previous research has established that key parts of this behavior, such as the approach, are visually guided, but it has remained unknown how mice would perform if given exposure to hunting without visual cues. Zona incerta, a potential key locus for the integration of external and internal information, has previously been shown in naïve mice to improve prey capture ability by increasing appetitive drive through integrating prey-related sensory information. It remains an open question if this region plays the same role in mice experienced with hunting in both the light and dark. Therefore, we examined the behavioral strategies used by mice with prior experience hunting in both the light and dark, as well as how these strategies are modulated by zona incerta. In experienced mice, we found that mice adjust their strategies to maintain similar success rates with and without visual cues, and that zona incerta further refines these strategies, specifically in the dark. This suggests that the role of zona incerta in these experienced mice is to maintain successful hunting by shifting behavioral strategies under ambiguous sensory conditions.

## Introduction

External sensory information and internal states combine to guide natural behaviors such as prey capture. While often thought of as a prey species, lab mice will readily chase, kill and eat insects after a few days of exposure^1^. This predatory hunting behavior can be visually-guided when mice hunt in the light, but much less is known about how they hunt in the dark, without visual cues^1,2^. As a primarily nocturnal species in the wild, and exclusively nocturnal in the lab, mice forage for insects predominantly at night^3,4^. Hunting in darkness is thus ethologically relevant for this species. Mice have evolved to use auditory, somatosensory, and olfactory information to hunt when vision is not available^4–8^.

Previous experiments restricting sensory information have revealed that prey capture is multimodal, with different cues dominating at different phases^1,2^. Without prior exposure to hunting in the dark, mice are significantly slower and show less directed approach angles than in the light, and earplugs exacerbate this effect, implicating auditory cues when vision is unavailable^1^. In the light, mice have access to visual, auditory, somatosensory, and olfactory information. In the dark with whiskers trimmed, mice lose visual and vibbrisal sources and must rely on auditory and olfactory information for much of the behavior, especially the approach phase which is often directed by vision^1^. While mice will use whatever sensory cues are available to hunt prey, how mice adjust their hunting tactics under these different sensory conditions remains unclear.

Sensory information must be unified with internal signals to guide successful behavior. The zona incerta (ZI) is a critical hub for integrating sensory information with internal state during appetitive behaviors. This diencephalic structure contains heterogeneous GABAergic populations positioned to gate information flow between cortex, thalamus, and downstream motor centers^9–12^. Consistent with its broad outputs, ZI participates in diverse functions including auditory cortical plasticity during threat learning, motor patterning through projections to the red nucleus, stress-induced grooming, anxiety-like phenotypes, and fear generalization^13–18^. Optogenetic experiments have also implicated ZI in prey capture. Activating medial ZI (ZIm) Vgat+ neurons projecting to periaqueductal grey (PAG) promotes hunting in naive mice by integrating prey-related sensory signals and inducing appetitive drive^19^. In these previous experiments, mice were exposed to crickets in their home cages, but naïve to hunting while tethered in the experimental arena, which has been previously shown to change aspects of prey capture such as the frequency of approach^20^. This population of ZI cells has also been established to target cells in PAG that are involved in prey capture motor patterning, food seeking, and food-related exploratory behavior^19,21,22^. These PAG-targeting cells in ZIm regulate the depth of investigation, driving a switch from “shallow” to “deep” exploration of novel objects^23^. Previous studies have also shown that experienced mice adapt to different strategies specifically during the pursuit phases of prey capture^24^.These findings suggest that with prior experience hunting, ZI may not initiate pursuit but rather modulates how animals engage with salient stimuli once detected.

We hypothesized that mouse hunting strategies would differ between light and darkness, and that these strategies depend in part on internal signals involving signaling in ZI. To test this, we recorded adult mice catching crickets in the light and dark with and without bilateral stimulation of ZIm inhibitory cells. We defined hunting in the dark with whiskers trimmed as “auditory-guided” prey capture, which does not exclude that mice have access to olfactory cues, but no longer have visual or vibrissal cues. We quantified the geometric relationships between predator and prey both as continuous variables and as discretized behavioral states. We found that mice were just as successful at prey capture in the dark as in the light, which they achieved by altering their movement strategies. Furthermore, we establish that in experienced hunters, ZI does not affect the initiation of prey capture behavior but instead increases persistence in engaging with the cricket once the behavior has begun. These results demonstrate how sensory input and internal state can combine to impact the structure of a natural behavior.

## Methods

### Animals

The transgenic model mice used for these experiments (Vgat-ChR2 mice) were generated by crossing Vgat-IRES-Cre mice (JAX 028862) with cdh23-ChR2 mice. The cdh23-ChR2 mice were in turn homozygous crosses of ChR2 mice (CAG-ChR2-eYFP, JAX 012569) with cdh23 rescue mice (JAX 018399). The cdh23 rescue mice were engineered with a single nucleotide substitution in the cdh23 gene to prevent age-related hearing loss in mice from the BL6 genetic background ^25^. These Vgat-ChR2 mice expressed channelrhodopsin (ChR2) in ZI Vgat+ cells. All experiments were conducted using male mice (n = 7) between 45 and 90 days of age (p45-90). All controls were within-animal no-laser trials; we did not include wildtype or GFP-only control mice in this study. Mice were housed in a 12-hour reversed light-dark cycle (lights off at 9:00 a.m.) in a temperature and humidity-controlled room. The animals were housed in standard cages (28.4 cm x 18.4 cm x 12.5cm) with ventilated cage lids and cedar bedding. In experiments using a viral strategy to deliver ChR2 (see below), Vgat-IRES-Cre mice (n=11) were first crossed to cdh23 rescue mice to generate a line with stable, high-frequency hearing.

### Optical fiber implants

Mice were implanted with bilateral optical fibers over the medial region of zona incerta (ZIm). Implantation coordinates were −1.80mm AP, −3.65mm DV, and +/− 1.50mm ML which were modified from Zhao et al., 2019 to better target ZIm following histological analysis in pilot cohorts of implants (Table S1). After surgery, mice were monitored for 5 days post-op and allowed to recover for at least a week before habituation procedures began.

### Viral injections

Vgat-Cre mice (n=11, 5M, 6F; p45-p90) were injected with a Cre-dependent ChR2 virus, pAAV-EF1a-double floxed-hChR2(H134R)-mCherry-WPRE-HGHpA (Addgene plasmid #20297). The injection coordinates and volume were based on Zhao et al., 2019 and targeting was validated histologically after behavioral experiments (Table S1). After viral injections, mice were given at least two weeks for sufficient expression of the virus in ZIm cell bodies before experiments were performed.

### Behavioral rig

All experiments were performed in a custom behavioral rig designed for freely moving mice to perform prey capture. We used a 60cm diameter, 30cm deep circular arena to provide a large area for mice to pursue the crickets. A high-resolution camera, IR LEDs for illumination, and white LEDs for illumination were suspended above the arena. Illumination was tested in the dark with the IR LEDs; narrow-band illumination peaked at 850nm and fell to background levels by 760nm, outside the visible range for mice. The camera was positioned to capture a top-down view of the entire arena and the behavior for later analysis. Video data and timing signals for aligning data streams were collected using Bonsai^26^. Optical fibers were suspended from the center of the behavioral rig along with the camera. Fibers were connected to a custom-built laser box, laser protocols were controlled by custom MATLAB software, and timing signals were recorded with OpenEphys^27^. Experiments were performed in a soundproof booth to isolate the animals from external auditory stimuli. Due to this, crickets were dropped into the arena at the beginning of a trial with externally controlled launchers. Remaining cricket detritus were cleaned in between each trial, and the arena was cleaned between mice.

### Habituation to the arena and crickets

To reduce anxiety, mice were first habituated for 3 days to the arena before prey capture experiments began. For habituation trials, mice were placed in the arena and connected to optic fibers first in the light and allowed to explore for 5-10 minutes. After habituation in the light, mice were allowed to explore the arena for 5-10 minutes in the dark. Habituation trial durations were 5 mins on the first day and were increased to 10 minutes over the 3 days. In addition to habituation to the arena, mice were also habituated to crickets before prey capture trials began. To habituate the mice to crickets, several crickets (Fluker Farms, 4-6 weeks old) were placed in their home cages overnight for the three nights before prey capture trials began.

### Whisker trimming

Whiskers were trimmed bilaterally after baseline light prey capture trials were completed before the dark trials (auditory-guided) began (see experimental timeline schematic). Mice were given the night to recover before starting prey capture the next day. Whisker length was monitored throughout the days of auditory-guided prey capture and whiskers were re-trimmed if the growth was more than 3 mm.

### Prey capture trials

#### Experienced hunters

Mice were not food restricted and had full access to food in their home cages. Before trials with optogenetic activation, mice performed several days of baseline prey capture trials, in order to increase their familiarity with hunting in the arena and while tethered. For baseline trials, mice were placed in the arena and attached to the optical fibers. Crickets were introduced by releasing them from one of two launchers. Mice were given 5 minutes to investigate, chase, and eat the cricket. If the mice didn’t catch the cricket within the 5 minute time limit, the cricket was removed, and the trial ended. If mice failed to catch the cricket on more than 5 consecutive trials, the mice were removed from the arena, and crickets were placed in their home cage overnight for further habituation. After 4 days of baseline prey capture trials with crickets in both the light and dark, prey capture trials with optogenetic activation began. Light and dark trials during both baseline and optogenetic activation were run in a block structure, with light days occurring before dark days (see experimental timeline schematic below). During optogenetic activation days, laser trials were pseudo-randomly interleaved with non-laser trials for each of the mice. For all mice, half of the trials for a day were laser, and the other half were non-laser. Control trials were within-animal laser OFF trials. Similar to baseline, each trial had a time limit of 5 minutes, and if mice failed to capture the cricket in that time, the cricket was removed, and a new trial started. If mice failed to capture in more than 5 consecutive trials, mice were removed from the arena, and no more trials were run for the day. Mice were run at the end of their light cycle or the beginning of their dark cycle, and were run in a pseudo-random order. Mice were excluded from group data analyses if they failed to capture crickets on more than 40% of trials (excluded n=1, final n=7).

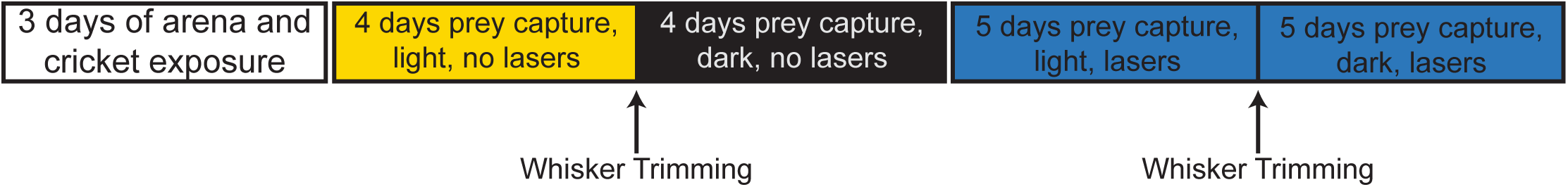

#### Naive hunters

Based on Zhao et al., 2019’s methods, we also included a group of Vgat-ChR2 mice (n=4M, p45-90) that were not first given baseline hunting in the light and dark before optogenetic activation. Instead, these mice were exposed to crickets in their home cages for three nights, and then optogenetic stimulation started on day 1 of prey capture in the light. Unlike the mice in Zhao et al., 2019, these mice were habituated to the arena before prey capture trials to reduce general anxiety-like phenotypes in the open field. These mice were considered “naive” hunters.

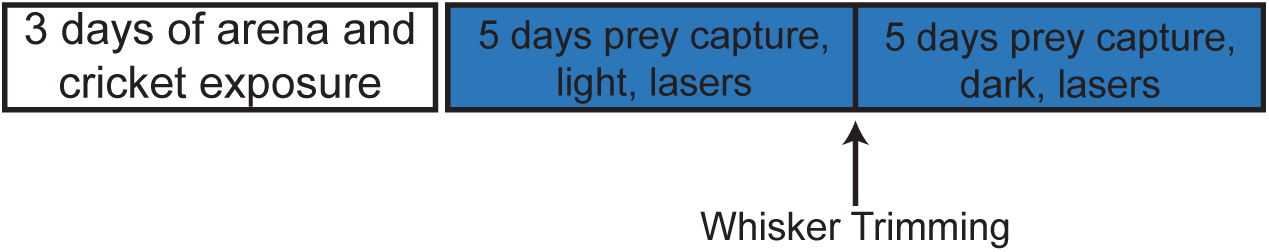

#### Optogenetic activation

ZIm Vgat+ cells were activated with a 473nm, 3.5 mW laser (111.4 mW/mm^2^)^28^. This laser power was chosen to be strong enough to activate ZI neurons while limiting the spread to any ChR2-expressing neurons outside ZI, based on previous validation experiments^19^. Activation protocols were modified from Zhao et al., 2019 with reductions to the laser power to limit spread to ChR2-expressing cells outside of ZI. The laser protocol was generated using custom Matlab software, and the timing of pulses was recorded using OpenEphys^27^. During laser trials, 10 ms pulses were delivered at 20 Hz for 2 minutes; laser stimulation started before the cricket drop, and stopped at the terminal capture (Fig. 1F). To ensure no illumination from the lasers was visible in the arena, ferrules were surrounded in an opaque heat shrink layer that was pushed down to the surface bottom of the fiber connection. Trials were stopped and not included in analysis if any light leakage was observed in the videos during recording. On each day that mice performed prey capture trials, half the trials were laser on, and half were laser off to allow for within-animal controls.

**Figure 1:**
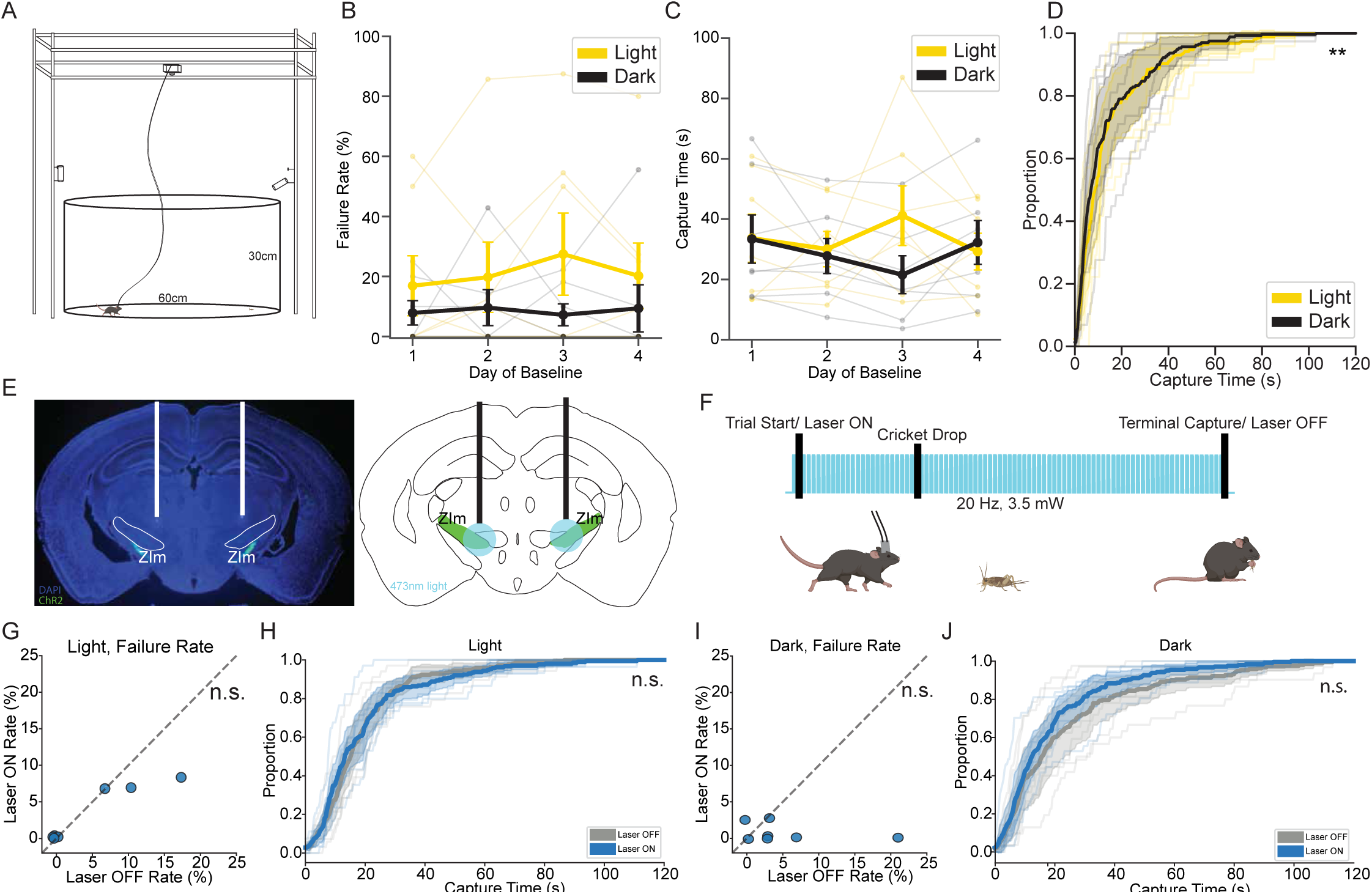
Activation of ZIm inhibitory cells does not decrease capture time or failure rate in experienced cricket hunters. A) Schematic of the behavioral arena. B) Failure rate did not differ significantly between light and dark across days of baseline (n=7; all days p>0.05, Wilcoxon Rank Sum, Holm Correction). C) Capture times did not vary significantly between the light and dark across days of baseline (n=7; all days p>0.05, Wilcoxon Rank Sum, Holm Correction). D) Mean capture times were slightly lower in the dark during baseline (n=7; p=0.004, K-S test). E) Example histology image and representation of ZIm and optic fiber location. F) Schematic of optogenetic stimulation parameters. G) Optogenetic activation did not significantly decrease failure rates (n=7; p=0.285, Wilcoxon Rank Sum) or H) capture times (n=7; p=0.284, K-S Test) in experienced mice in the light. I) Optogenetic activation did not significantly decrease failure rates (n=7; p=0.075, Wilcoxon Rank Sum) or J) capture times (n=7; p=0.248, K-S Test) in experienced mice in the dark.

In additional cohorts of experienced Vgat-Cre hunters (Fig. S6) with viral expression of ChR2, 10ms pulses were delivered at 20 Hz for 2 minutes at 6.3mW, using a higher power laser because there was no chance of off-target ChR2 expression, unlike in the transgenic mice.

#### Histology and fiber validation

After experiments were completed, mice were perfused with 4% PFA, the brains were removed, and then post-fixed in 4% PFA overnight. After post-fixation, 30 um slices were taken on a vibratome. Slices were mounted using DAPI Fluoromount to visualize cell bodies. Images were taken to verify the expression of ChR2 in Vgat+ cells as well as the location of optic fibers. For Vgat-Cre mice expressing the ChR2 virus, signal from the mCherry tag was amplified using an anti-dsRed primary antibody (1:500, Takara Bio Living Colors® DsRed Polyclonal Antibody) and AlexaFluor-488 secondary antibody (1:1000). Mice (excluded n=2, final n=7) were excluded from the group data analysis and total reported n if the fiber location was not directly superficial to ZIm.

#### Behavioral data analysis

To analyze the behavior of mice during prey capture, we trained a multi-animal DeepLabCut (maDLC)^29^ model to track both the mouse and the cricket positions throughout the entirety of the trial. We used custom Python code to extract the positions of the mouse and cricket from maDLC output, and then calculated geometric variables to describe the behavior, i.e. range, azimuth, mouse speed and cricket speed. These variables were key to calculating metrics of prey capture success such as time to capture, latency to attack and capture probability. Approaches (azimuth less than 30 degrees and speed greater than 5 cm/s) and intercepts (approach periods ending in range less than 5 cm) were modified from thresholds reported in Hoy et al., 2016. We used conjunctions of different geometric variables to identify discrete events (occurring at a single point in time) and states (which were extended in time) within the prey capture behavior. We defined 6 states (pause, wander, stalk, follow, chase, hot pursuit) using the following criteria: pause: mouse speed < 2 cm/s; wander: mouse speed 2-15 cm/s, range > 20cm, azimuth >30 deg; stalk: mouse speed 2-15 cm/s, range 5-20 cm, azimuth > 30 deg; follow: mouse speed 5-15 cm/s, range 5-20 cm, azimuth < 30 deg; chase: mouse speed 15-25 cm/s, range 5-20 cm, azimuth < 30 deg; hot pursuit: mouse speed 25-100 cm/s, range 5-20 cm, azimuth < 30 deg. We defined 7 discrete behavioral events from geometric variables using the following criteria: cricket jump: peaks in the cricket speed; jump loss: cricket jumps followed by an increase in range and azimuth; range min: local minimum in the range; range min follow: a range min followed by maintaining low range and azimuth; range min loss: a range min followed by increased range and azimuth; failed approach: an approach period ending in range >5 cm; intercept (contact): approach period ending in range < 5 cm. We developed and refined this set of states and events, and the combinations of geometries used to define them, based on experimenter observations of prey capture behavior across the full span of testing. Partial transition probability matrices were generated from behavioral states. Since our state labeling did not assign a state to every frame of data, unlabeled frames were added to an “unclassified” state that was excluded from transitions.

### Statistical tests

Because datasets were not normally distributed, we used Wilcoxon paired rank-sums for failure rates. Significance in capture times was calculated using the Kolmogorov-Smirnov (K-S) test. For the drop distance interactions, a quantile regression on medians and mixed linear model on means was used to test for an interaction between drop distance and laser condition. Mixed linear models that included mouse as a random effect were run to test for differences in the trial median geometries, and pre-contact median geometries were tested with Wilcoxon paired rank-sums. To assess significance in changes to geometric distributions, we calculated the Wasserstein Earth Mover’s Distance (EMD) as a measure of how much energy it would require to move one distribution into the other. Reported sigma values represent the number of standard deviations our EMD was from the null EMD calculated from 1000 permuted shuffled null distributions (Fig. S2, 5); sigma>6 corresponds to p<0.001. For behavioral states, negative-binomial GLMs were used to test for differences in the per trial counts. To create a null partial TPM we shuffled trial labels within each mouse to preserve the real transition structures but change the laser labels and permuted this shuffle 10,000 times. We compared the observed transition structure to the null and ran a false discovery rate correction for multiple comparisons. We calculated observed EMD between the joint probabilities of the given conditions from a 10×10 histogram, shuffled the frame labels within mouse within day and permuted this 1000 times to generate null joint probability distributions. Sigma values were calculated as number of standard deviations from the null EMD.

## Results

### Mice rapidly learn to pursue and catch crickets, even under ambiguous sensory conditions

Given that mice are predominantly a nocturnal species, and often hunt or forage in low light or complete darkness, we wondered how mice would perform at auditory-guided prey capture given several days of exposure after they had become proficient at prey capture in the light ^4,5^. To investigate this, we allowed mice four days of exposure to catching crickets with all sensory information present (visual, auditory, somatosensory and olfactory) in a large 60cm diameter arena (Fig. 1A). After 4 days of hunting under these conditions, we then removed visual and vibrissal cues by having mice hunt in complete darkness with whiskers removed. We found that as early as on day 1 of auditory-guided prey capture, mice were already catching crickets as successfully and as quickly as in the visually-guided paradigm (n=7; p>0.05, Wilcoxon Rank Sum, Holm Correction; Fig. 1B-C). Pooled across all days, their capture times were significantly lower in the auditory-guided baseline period than in the light baseline hunting period (n=7; p<0.01, K-S Test; Fig 1D). The differences between capture times in the light and dark baseline likely reflect additional learning and refinement of prey capture, since our experimental design meant that mice had more hunting experience by the time they were hunting in the dark (Fig. 1B-D). Nevertheless, these results demonstrate that given previous exposure to catching in the dark, mice can perform auditory-guided prey capture and effectively hunt crickets in the absence of visual cues.

### Activation of ZIm inhibitory cells does not affect capture times or failure rates in experienced mice

Medial ZI has been shown to promote visually-guided prey capture in naive mice, but given how quickly mice become experienced hunters in both the light and the dark (Fig. 1), we wondered whether ZI has a similar role in experienced mice. We implanted male Vgat-ChR2 transgenic mice (n=7) with optic fibers over ZIm to activate Vgat+ inhibitory cells (Fig. 1E)^19^. These mice were exposed to crickets in their home cages and allowed to become experienced hunters in both the light and dark before any trials with optogenetic activation began. A 473 nm laser calibrated for 3.5 mW intensity was pulsed at 20 Hz for the first 120 seconds of trials; optogenetic stimulation started 5 seconds before the cricket dropped and was stopped at the terminal capture (Fig. 1F). Activation trials were pseudo-randomly interleaved with non-activation trials so that every day mice hunted both with and without optogenetic activation of ZIm.

We found that when mice hunted in the light and dark, there was little effect of optogenetic activation of ZIm on overall success metrics. With and without visual cues, optogenetic activation did not significantly reduce capture times or failure rates (n=7; p>0.05, capture times: K-S Test, failure rates: Wilcoxon Rank Sum; Fig. 1G-J). We wondered if the difference in effects between our experienced mice and previous findings in naïve (viral) mice could be due either to hunting experience or to the specificity of the expression of ChR2. To test this, we collected prey capture data from experienced mice with ChR2 restricted to ZIm and from naïve Vgat-ChR2 transgenic mice. Neither of these groups showed significantly shorter capture times or lower failure rates with ZIm activation (Fig. S6-7). These results suggest that the difference between our experienced mice that showed no effect, and previous reports of ZIm effects in naïve (viral) mice cannot be explained by the specificity of ChR2 expression alone^19^.

Because of the dynamic and variable nature of prey capture, single-number metrics such as capture time offer an incomplete picture of this behavior. One source of variability is whether the mouse quickly detects and begins chasing the cricket, which in our experiments depended in part on the random location at which the cricket is dropped into the arena. To control for this source of variability, we partitioned the data into quartiles based on the initial distance between mouse and cricket at the drop. This partitioning revealed a significant interaction between drop distance and laser, indicating that the laser decreased capture times in the shortest drop quartile (n=7; p<0.01, Quantile Regression; Fig. S1 B-C). This is consistent with previous reports that ZIm activation decreased capture times in an arena as small as our shortest drop distance quartile^19^. Thus, in experienced mice, we found that ZIm activation did not shorten capture times across the board, but it did shorten them specifically on trials in which the cricket was dropped closer to the mouse – that is, on trials that consisted mostly of pursuit. This suggests that ZIm may enhance prey capture ability only after mice are already engaged in chasing the cricket.

### Geometric relationship between mouse and cricket differs depending on sensory information available

Prey capture inherently involves the dynamic interactions between two individuals. To quantify the geometry of these interactions, we calculated range, azimuth, mouse speed and cricket speed from the positional information we extracted from DeepLabCut tracking (Fig. 2A)^13^. We found that median range in the dark was significantly lower than in the light, whereas median speed was significantly higher in dark than in light (n=7, range: p<0.001, speed: p<0.001, mixedLM). Thus, mice moved faster and spend more time closer to the cricket in the dark than in the light (Fig. 2B-D). These differences suggest that without visual cues, mice may adopt a different strategy to maintain their successful hunting performance.

**Figure 2:**
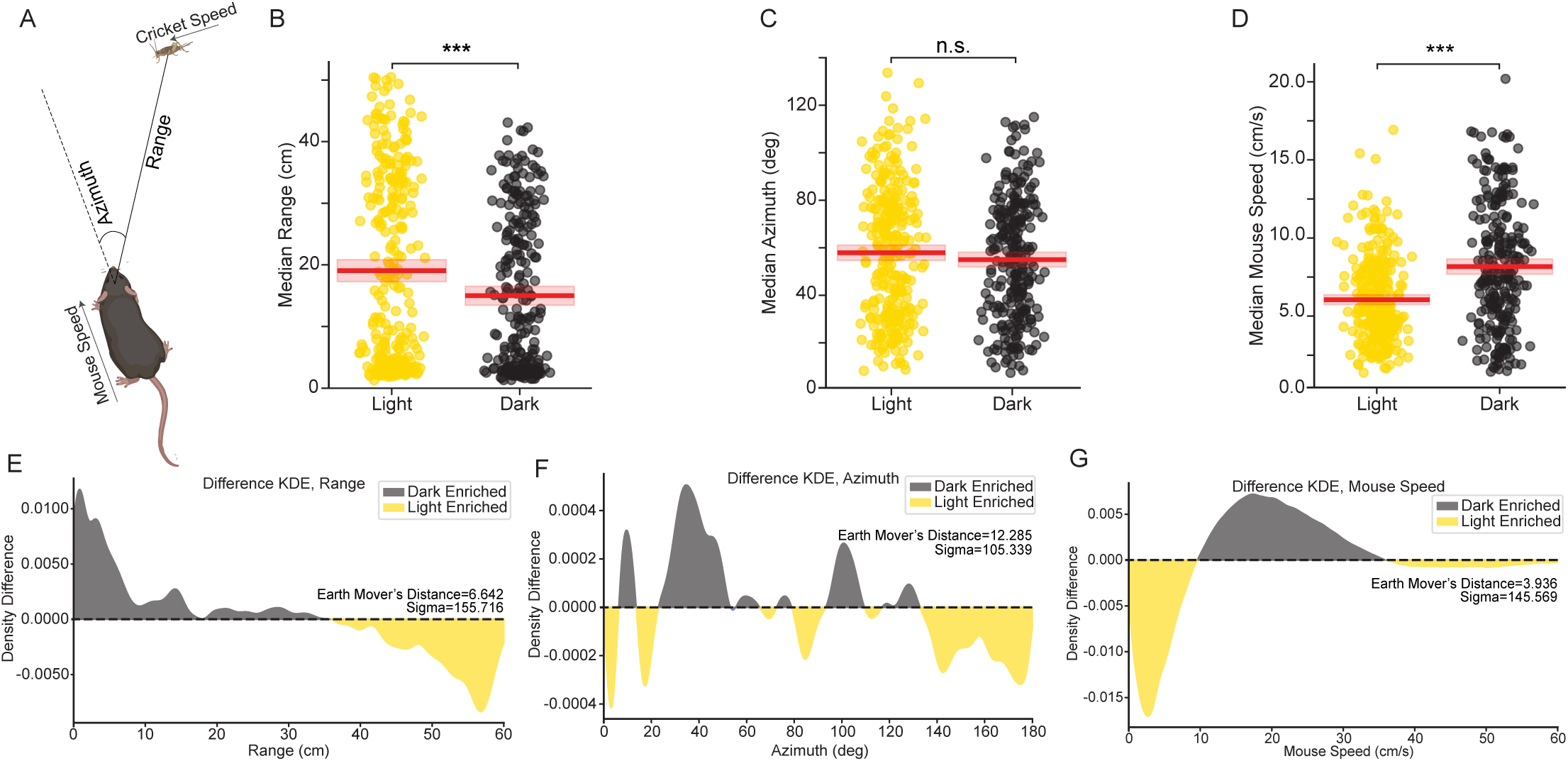
Geometric variables reveal differences in mouse-cricket interactions under different sensory conditions. A) Schematic of the different calculated geometric variables. B) Median range was significantly lower in the dark than in the light (n=7; p=1.467e-6, mixedLM). C) Median azimuth did not significantly differ between light and dark (n=7; p=0.071, mixedLM). D) Median speed was significantly higher in the dark than in the light (n=7; p=6.638e-16, mixedLM). E) Distribution of ranges revealed significantly lower ranges in the dark (n=7; sigma=155.716, p<0.001, EMD Permutation Test). F) Distribution of azimuths shifted significantly in light compared to dark (n=7; sigma=105.339, p<0.001, EMD Permutation Test). G) Dark significantly increased medium speeds compared to the light (n=7; sigma=145.569, p<0.001, EMD Permutation Test).

Because the geometric variables change continuously throughout each trial, the trial medians in Fig. 2B-D do not reveal the full picture of behavioral dynamics. Therefore, we visualized the distributions of these variables using kernel density estimation (KDE), which represents the full relationship between mouse and cricket throughout the trial. To quantify the differences between geometric distributions, we calculated the Earth Mover’s Distance (EMD) between the two distributions. The difference in the distribution of ranges indicates an increase of shorter ranges in the dark compared to light (n=7; p<0.001, EMD Permutation Test; Fig. 2E, Fig. S2). This change in the range distribution demonstrates that in the dark, mice maintain closer contact with the cricket to successfully catch it. In the difference plot of azimuth between mouse and cricket, the difference between light and dark was more complex, but nonetheless highly significant (n=7; p<0.001, EMD Permutation Test; Fig. 2F, Fig. S2). The difference in the distribution for the mouse speeds indicated that the increase in median speed was not due to an increase of the highest speeds, but rather an increase in intermediate speeds along with a down-regulation of the slowest and fastest speeds (n=7; p<0.001, EMD Permutation Test; Fig. 2G, Fig. S2). During auditory-guided prey capture, mice only receive auditory information about cricket location when the cricket is moving. Thus, moving faster in the dark might reflect a strategy to “flush” the cricket, that is, to cause it to move in order to obtain auditory information about its location.

In addition to these differences that emerged from the entire behavior taken as a whole, we also wondered whether geometric variables might differ at key timepoints during prey capture, such as the moment of contact between mouse and cricket, or the initiation of an approach. Aligned to intercept events, we found that after four days of baseline catching in the dark, mice had higher median azimuths preceding contact than in the light (n=7; p<0.05, mixedLM; Fig. S3A-B). Additionally, the mean azimuth at the start of the first approach to the cricket was significantly lower in the light than in the dark (n=7; p<0.05, Wilcoxon Rank Sum; Fig. S3C). The nature of these differences suggests that to be as successful in the dark, mice employ different tactics than in the light. That is, mice specifically stay closer to crickets and move faster during prey capture in the dark, while at the start of approaches they are less tightly oriented to the cricket.

### Mice more closely engage with the cricket during optogenetic activation of ZIm inhibitory cells

These findings reveal that the relationship between mouse and cricket, which is key for success in hunting, is modulated by sensory condition. We therefore asked if this relationship could further be shifted by optogenetic activation of ZIm. Given the role of ZI in depth of investigation in other tasks, we wondered if activation of these cells might shift mice toward more engagement with the cricket, leading to more efficient prey capture strategies^23^. We found that the distributions of geometric variables revealed details about the underlying structure of this behavior, as well as shifts driven by optogenetic activation of ZIm cells. ZIm optogenetic activation produced highly significant shifts in range in both light and dark prey capture, which difference plots revealed to be an up-regulation of closer ranges between the mouse and cricket (n=7; p<0.001, EMD Permutation Test; Fig. 3A, E, Fig. S5C-D). ZIm activation also produced significant shifts towards smaller azimuths in both light and dark prey capture (n=7; p<0.001, EMD Permutation Test; Fig. 3B, E, S5A-B). In the light, mice shifted towards slower speeds, whereas in the dark mice showed an up-regulation of both the slowest and fastest speeds while the speeds in the middle were depleted (n=7; p<0.001, EMD Permutation Test; Fig. 3C, F, S5E-F). These shifts suggest that even without changes in global success metrics such as capture time, ZI activation causes mice to employ different tactics to stay more persistent at maintaining closer contact with the cricket once engaged in hunting.

**Figure 3:**
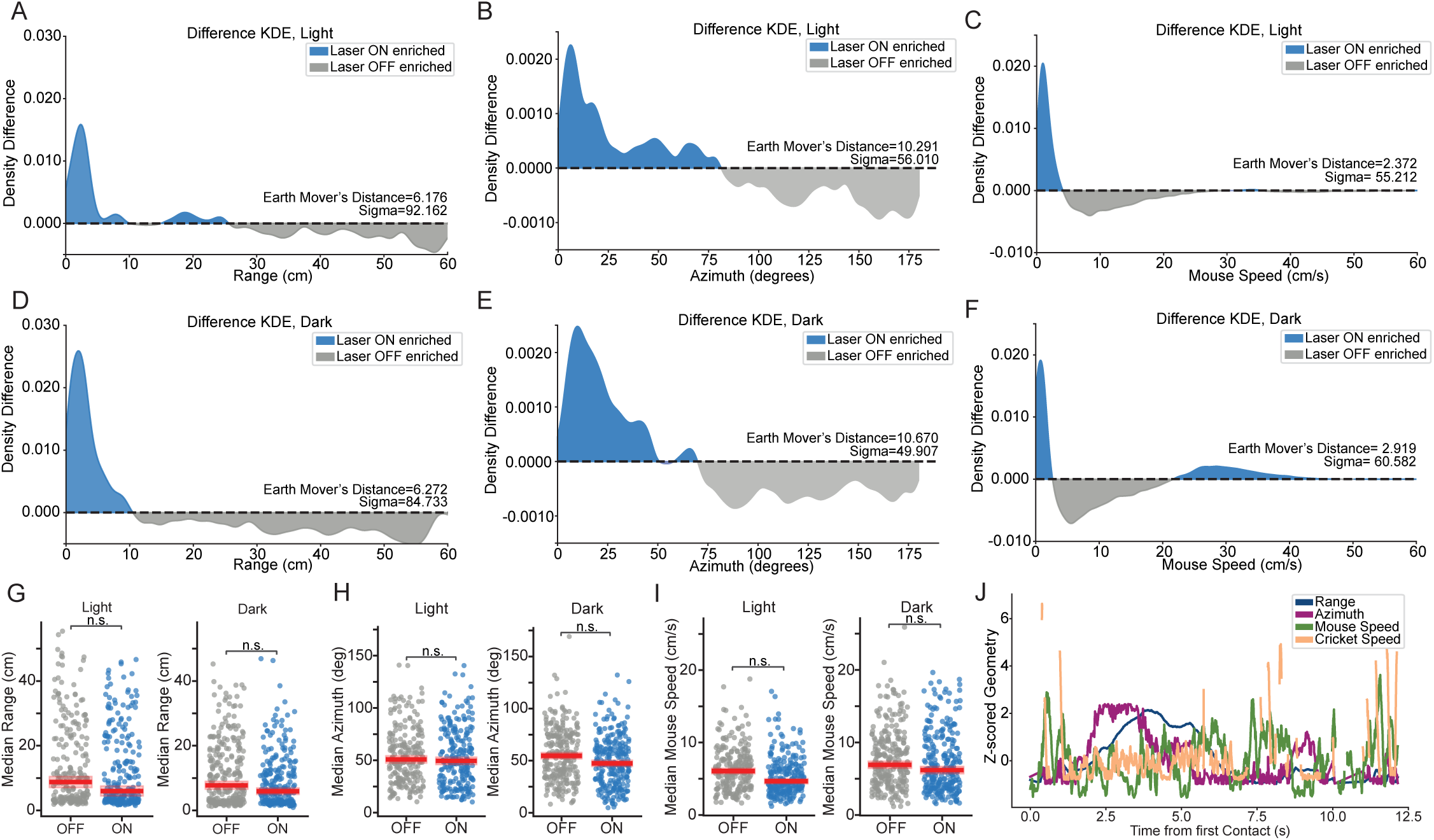
Activation of ZIm inhibitory cells shifts distributions of geometric variables in both light and dark. A) Optogenetic activation of ZIm shifted distributions towards shorter ranges (n=7; sigma=92.162, p<0.001, EMD Permutation Test), B) smaller azimuths (n=7; sigma=56.010, p<0.001, EMD Permutation Test), and C) slower speeds (n=7; sigma=55.212, p<0.001, EMD Permutation Test) in the light. D) Activation of ZIm shifted distributions towards shorter ranges (n=7; sigma=84.733, p<0.001, EMD Permutation Test), E) smaller azimuths (n=7; sigma=49.907, p<0.001, EMD Permutation Test), and F) both the slowest and fastest speeds (n=7; sigma=60.582, p<0.001, EMD Permutation Test) in the dark. G) Activation of ZIm did not significantly change median range (n=7; light: p=0.922; dark: p=0.300, mixedLM) H) azimuth (n=7; light: p=0.980; dark: p=0.207, mixedLM) or I) speed (n=7; light: p=0.124; dark: p=0.782, mixedLM) in the light or dark. J) Example z-scored geometries for one trial indicated the variability of these measures.

When we examined the median range, azimuth and mouse speed for each trial, we found that there were no significant effects of ZIm activation in either the light or dark (n=7; p>0.05, mixedLM; Fig. 3G-I). Given the variability inherent in this behavior, it is not surprising that a manipulation may not result in large shifts in single summary metrics per trial. Prey capture is a complex and highly dynamic behavior, and all geometric variables show large fluctuations throughout an individual trial (Fig. 3J). Unlike the difference between light and dark trials during the baseline period, in these experienced mice stimulation of ZIm did not introduce a large change to the system (Fig 3G-J).

### Mice utilize different strategies when ZIm is activated, particularly during periods of high engagement in the dark

Given our findings that mice were significantly closer and more tightly oriented towards the cricket when ZIm cells were optogenetically activated, we wondered what the relationship between mouse and cricket would look like during key moments of the behavior. To answer this question, we looked at the azimuth and speed preceding and following intercepts, and found that in both the light and dark, mice maintained smaller azimuths both before and after intercepts during optogenetic activation (n=7; light: p<0.05, dark: p<0.05, Wilcoxon Rank Sum; Fig. 4A-D). In addition, activation of ZIm inhibitory cells led to higher speeds before intercepts in the dark, while speeds did not significantly change in the light (n=7; light: p>0.05, dark: p<0.05, Wilcoxon Rank Sum; Fig. 4E-H). Unlike the baseline differences between light and dark, we found that activation of ZIm did not change the azimuth at the start of the first approach or at the cricket drop (n=7; p>0.05, Wilcoxon Rank Sum; Fig. S4). Decreased median azimuth and increased speed before intercepts indicate that during key periods of hunting, i.e. immediately before a contact with the prey, mice have shifted their tactics to a more direct and faster approach. This shift in tactics further indicates that mice may utilize different strategies during periods of high engagement with the cricket during optogenetic activation of ZIm cells.

**Figure 4:**
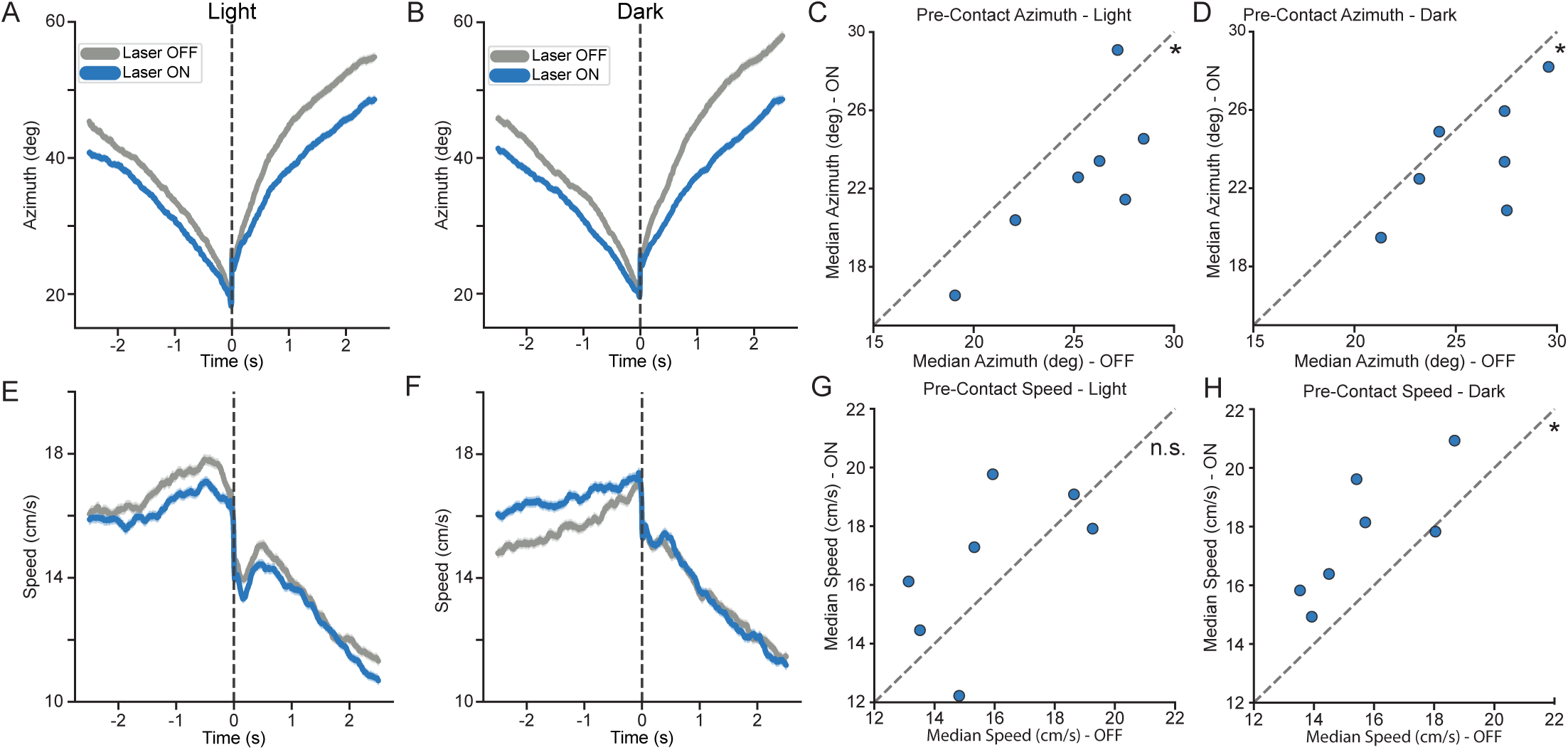
ZIm activation alters pre-contact azimuth and speed in the dark. A) Time series of pre-contact azimuth was lower in the light and B) dark. C) Median pre-contact azimuth was significantly lower in the light (n=7; p=0.047, Wilcoxon Rank sSum) and D) dark (n=7; p=0.031, Wilcoxon Rank Sum). E) Time series of pre-contact mouse speed was lower in the light, and F) higher in the dark. G) Median pre-contact speed did not significantly change in the light (n=7; p=0.375, Wilcoxon Rank Sum), but H) did in the dark (n=7; p=0.031, Wilcoxon Rank Sum).

### Geometrically defined state analysis reveals the underlying structure of prey capture behavior

The complexity of prey capture and the continuously varying interactions between mouse and cricket led us to define different behavioral states and events based on combinations of geometric relationships. We defined 6 states ranging from high-speed, tight-angle and close-range (more “pursuit-like”) to slow, long-range and wide-angle (more “search-like”) (Fig. 5A-C). These geometrically defined states, ordered from most-engaged to least-engaged, based on the center of the speed window, were hot pursuit, chase, following, stalk, wander, and pause.

**Figure 5:**
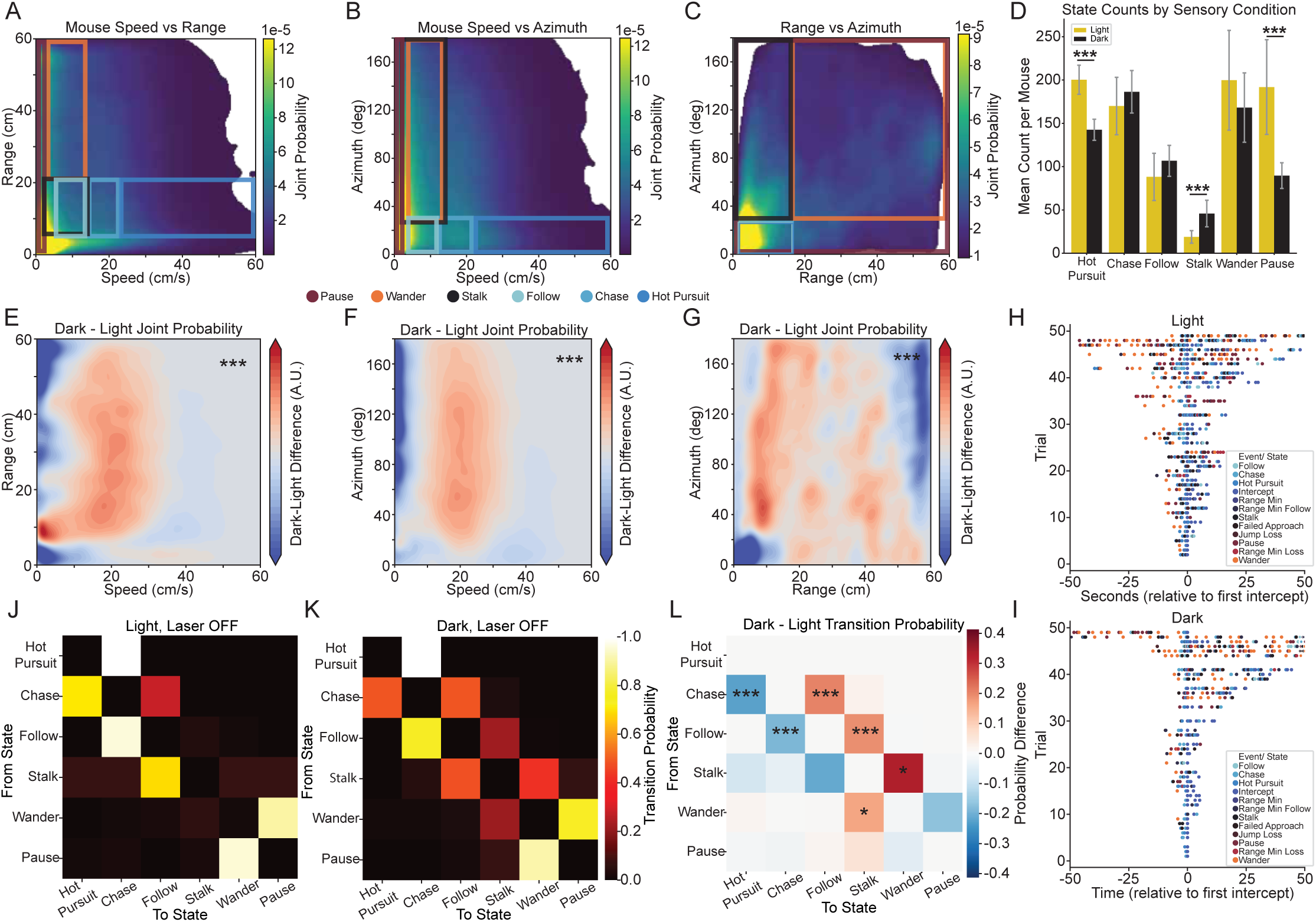
Joint probabilities and combinations of geometries can be used to label different behavioral states and events. A-C) Boundaries drawn on joint probabilities indicate cutoffs used to isolate discrete behavioral states. D) Mean counts of hot pursuit (p<0.001, Negative Binomial GLM), stalk (p<0.001, negative binomial) and pause (p<0.001, Negative Binomial GLM) are significantly different in light and dark, all other state counts did not change significantly (p>0.05, Negative Binomial GLM). E-G) Difference plots of joint probabilities of geometric variables revealed shifts under different sensory conditions (n=7; Range-Speed: Sigma=442.975, p<0.001, Azimuth-Speed: Sigma=242.564, p<0.001, Azimuth-Range: Sigma=466.737, p<0.001; EMD Permutation Test). H-I) Sequences of behavioral states suggested the structure of prey capture behavior does not support a canonical model of search phases followed by chase phases. J-L) Partial transition probability matrices indicate differences in the baseline structure of this behavior under different sensory conditions (n=7; *** p<0.001, *p<0.05, Permutation Test, FDR Correction).

Most of these states occurred at broadly similar rates, but certain states were more common under different sensory conditions (Fig. 5D). Specifically, both hot pursuit and pause states were more common in the light, whereas stalk was more common in the dark (n=7; all p<0.001, Negative Binomial GLM; Fig. 5D). In addition to defining behavioral states (which are extended in time), we used similar logic to isolate discrete behavioral events (which occur at a moment in time) such as cricket jumps, intercepts, and failed approaches. We also used the difference plots of joint probabilities of geometric variables to examine differences in hunting tactics in the light and dark. Like the distributions of single geometric variables, these represent the full relationship between the mouse and cricket throughout each trial. The joint probabilities revealed that in the dark, mice were moving slower when closer to the cricket, which is consistent with the down-regulation in states such as hot pursuit and the up-regulation of stalks (n=7; p<0.001, EMD Permutation Test; Fig. 5E). Overall, all the joint probabilities revealed a down-regulation of the closest range-azimuth and range-speed combinations, and a slight up-regulation of the intermediate azimuth-speed, range-speed and range-azimuth combinations in the dark (n=7; all p<0.001, EMD Permutation Test; Fig. 5E-G). This suggests that mice hunting in darkness rely more heavily on a stalking strategy using intermediate speeds, azimuths, and ranges.

We wondered whether prey capture would follow a canonical sequence of states and events, such as search generally leading to pursuit, as has been described for prey capture in other species such as echolocating bats. To investigate this, we aligned behavioral states to the first intercept in each trial and examined the relative timing of different states. We found that state and event sequences were highly variable across trials, with no strong evidence for any “search” or “pursuit” phases of the behavior in these experienced mice (Fig. 5H-I). Given this variability, we next turned to a probabilistic approach to characterize state sequences based on their transition probabilities.

### State transitions and event interactions reveal underlying structure of this complex behavior

To examine the structure of transitions between different behavioral states, we constructed partial transition probability matrices from the different behavioral states. These partial TPMs revealed in both light and dark prey capture there are strong tendencies for states to transition into their neighboring states when ordered from most to least engaged with the cricket (based on the speed window centers used to define them, Fig. 5J-K). There were modestly different transition structures in the light compared to the dark, with transitions in the dark having down-regulated transitions between the more engaged states, and more transitions between the less engaged states (n=7; ***p<0.001, *p<0.05, Permutation Test, FDR Correction; Fig. 5J-L, Table S2). The TPM was strongly banded, indicating a continuum from the fastest to slowest states, without strong clustering into categories of search-like or pursuit-like states. This can be seen from the near-absence of off-diagonal structure (Fig. 5J-K). Thus, an intuitive search-pursuit model of this predator-prey interaction is oversimplified. Rather than forming distinct, well-separated search and pursuit behavioral modes, these findings suggest a single continuous axis of pursuit engagement. The structure of transitions provides evidence for a continuum of behavioral phases determined by proximity to the cricket^7^. The differences in transitions between light and dark likely reflects the viability of different search strategies given different sensory information. Without visual cues, stalking becomes a much more valuable strategy, because more movement from the mouse is likely to promote more movement from the cricket, generating more auditory stimuli for the mice to track down.

### Activation of ZIm inhibitory cells promotes more engaged states while hunting in the dark

Having established a framework for identifying behavioral states and events from our geometric variables, we next asked how these states and transitions were affected by ZIm stimulation. Without visual cues, the less-engaged states (stalk, wander) were less common during optogenetic activation, as were the slowest of the engaged states (follow) (n=7, stalk, wander: p<0.001, follow: p<0.01, Negative Binomial GLM; Fig. 6A). In contrast, when mice had visual cues, activation of ZIm had no effect on state counts (n=7; all p>0.05, Negative Binomial GLM; Fig. 6B). We also wondered if activation of these inhibitory cells would drive changes to the transition structure. To answer this question, we used partial TPMs and computed the difference between laser on and off. We found that when mice were hunting without visual cues (auditory-guided), activation of ZIm shifted the transitions into “engaged” state types; most transitions tended to be towards a state’s neighboring, more engaged state rather than into the more disengaged state (n=7; **p<0.01, *p<0.05, Permutation Test, FDR Correction; Fig. 6C, Table S4). However, in the light, none of the transitions were significantly modulated by optogenetic activation of ZIm (n=7, all p>0.05, Permutation Test, FDR Correction; Fig. S8, Table S3). Both the transitions and state counts revealed that with optogenetic activation, mice spent more time closely engaging with the cricket, suggesting one possible effect of these cells is to increase persistence during prey capture, particularly under more difficult or ambiguous sensory conditions.

**Figure 6:**
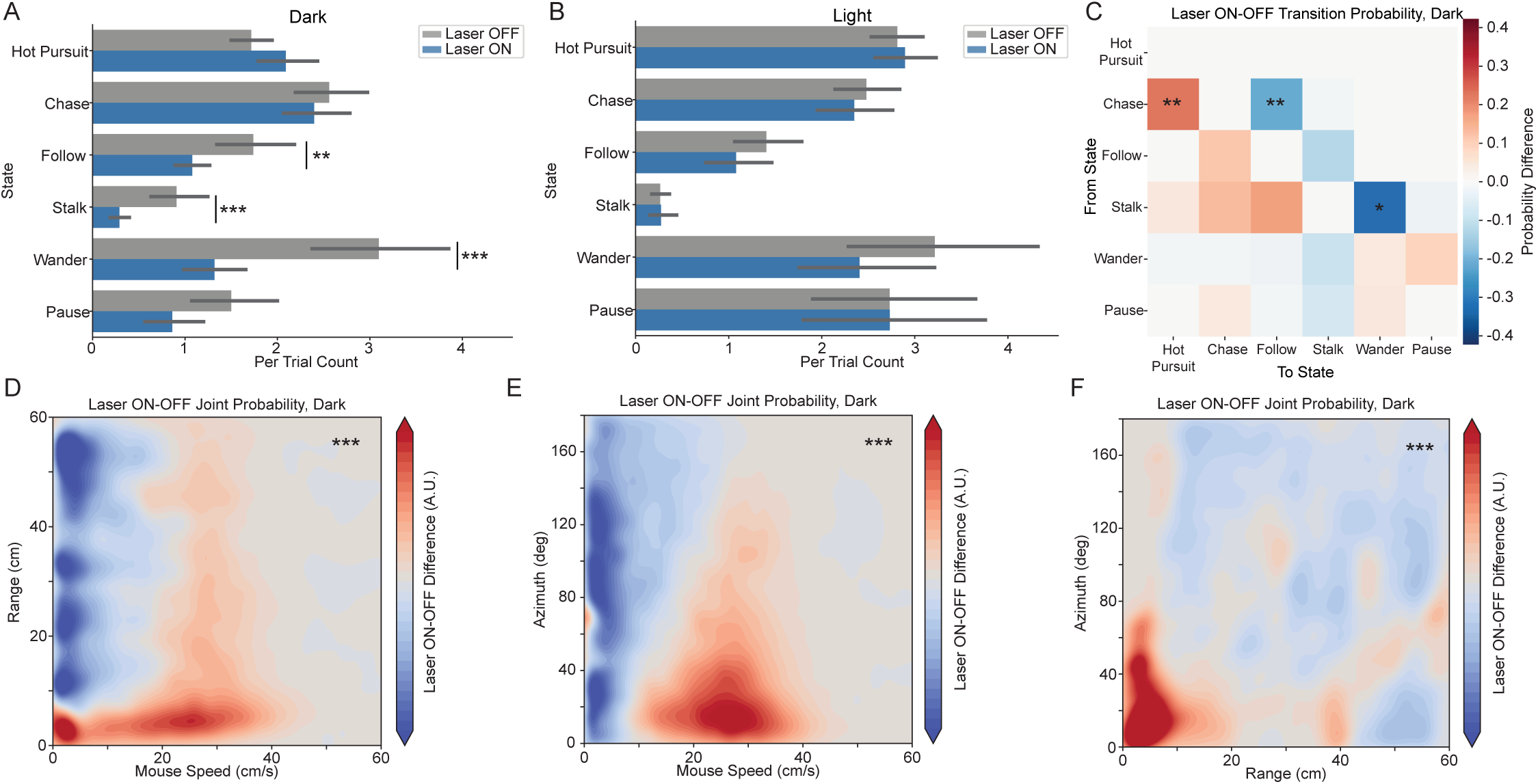
Activation of ZIm inhibitory cells shifts the structure of prey capture behavior in the dark but not in the light. A) Optogenetic stimulation decreased the occurrences of follow (p=0.003, Negative Binomial GLM), stalk (p<0.001, Negative Binomial GLM) and wander (p<0.001, Negative Binomial GLM), but not other states (all p>0.05, Negative Binomial GLM) in the dark. B) Counts of the different behavioral states did not significantly change in the light (all p>0.05, Negative Binomial GLM). C) Partial transition probability matrices revealed a shift towards more transitions in and out of the coupled/ engaged states (n=7; ** p<0.01, *p<0.05, Permutation Test, FDR Correction). D-F) Difference plots showed activation of these cells shifted the joint probabilities of geometric variables into more closely coupled/ engaged distributions, i.e. higher speeds, lower ranges and azimuths (n=7; Range-Speed: Sigma=313.357, p<0.001, Azimuth-Speed: Sigma=181.246, p<0.001, Azimuth-Range: Sigma=361.072, p<0.001; EMD Permutation Test).

Given the ZIm-evoked shifts in geometric variables, and in states that are defined by combinations of geometric variables, we wondered if these changes might also be reflected in the joint probabilities. We found that ZIm activation shifted the joint probability of speed and azimuth towards higher speeds and lower azimuths. Speed and range joint probabilities were also shifted towards higher speeds and lower ranges, and the azimuth and range joint probability moved to low ranges and azimuths (n=7; all p<0.001, Permutation Test; Fig. 6D-F). Taken together with the changes to state counts and transitions, these effects suggest that ZIm does not simply cause mice to chase more, but rather changes the nature of their interactions with the cricket, to stay more in a more persistent regime.

## Discussion

Animals must adjust active sensing strategies to their internal state and available sensory information when executing natural behaviors like prey capture. As a primarily nocturnal species, wild mice hunt in darkness more often than in light, yet laboratory studies have focused on visually-guided hunting^1,4^. In this study, we show that mice maintain prey capture success in darkness by altering their movement strategies. Further, we establish that activating inhibitory neurons in the zona incerta (ZIm), a region implicated in integrating sensory information with internal state, also shifts hunting strategies toward closer, more persistent engagement without changing overall capture success. Notably, many of the behavioral shifts we report emerged not in single-value metrics like mean capture time, but in the full distributions of predator-prey geometry. This suggests that mice adjust how they actively sample sensory information through changes in behavioral strategies, such as moving faster to flush the cricket, to achieve similar levels of overall performance. This flexibility may allow mice to maintain robust prey capture performance even when sensory conditions and internal state often change substantially.

Hunting in darkness involves strategic adjustments that mice adopt when hunting without visual information. After only a few days of experience, mice hunting in darkness with whiskers trimmed (auditory-guided prey capture) maintained closer ranges to prey, moved at higher intermediate speeds, and relied more heavily on stalking behaviors compared to hunting in the light. These findings extend previous work showing that vision guides long-range approach during prey capture by demonstrating that audition becomes sufficient when mice gain experience with the task in the dark ^1^. While olfactory cues likely contribute to motivational state, they are unlikely to provide precise spatial information about cricket location^1,6,7^. Together, these results indicate that mice do not simply perform worse in the dark but instead deploy distinct strategies optimized for auditory-guided pursuit to hunt just as successfully as in the light.

Activating ZIm inhibitory cells also alters prey capture strategies in experienced mice by increasing persistence once engagement has begun, rather than initiating pursuit. In our experienced hunters, ZIm activation did not affect capture time or failure rates, but instead shifted mice toward closer, more sustained engagement with the cricket after pursuit started. This pattern differs from previous research showing that activating ZIm promotes hunting in naive mice by inducing appetitive drive^19^. One explanation for this difference is that ZIm cells are also involved in depth of investigation of novel objects^23^. Prior experience with hunting decreases the novelty of crickets in the context of the experimental arena, reducing the contribution of novelty-seeking mechanisms. Rather than modulating appetitive drive, our results suggest ZIm sustains engagement with salient stimuli once detected, particularly when sensory conditions are ambiguous. These findings fit with research in other species that showed internal state as key in shifting behavioral states during a hunting task^30^.This indicates that ZIm integrates sensory information with internal state to maintain goal-directed pursuit, not simply to initiate it.

The effects of ZIm activation on hunting strategies were specific to prey capture in the dark. During auditory-guided hunting, ZIm stimulation shifted behavioral state transitions toward more engaged states and increased the probability of maintaining close contact with prey. In contrast, these effects were absent when mice hunted with visual cues available. The sensory environment in the light and dark is fundamentally different, likely leading to more uncertainty when hunting without visual cues. Prey capture behavior is driven by the interplay between approach and avoidance behaviors, mediated in part by cells in lateral hypothalamus and periaqueductal gray^31^. ZI sends projections to both regions, and efferent projections from this region have been implicated in defensive flight behaviors as well as anxiety phenotypes^7,32,33^. This circuitry for motivational control of hunting may enable ZIm to modulate engagement with salient stimuli when sensory evidence is limited, thereby counteracting a bias towards avoidance in ambiguous conditions. This is consistent with our observation that ZIm effects emerge specifically during dark hunting conditions.

Given its many functions involved in sensory integration and internal state processes, zona incerta is a compelling region to investigate during complex, goal-directed behaviors. Our findings expand the role of this region beyond appetitive drive to include persistence of engagement when sensory evidence is limited. Just as different distributions can share the same mean, different neural mechanisms can produce the same capture success. ZIm activation did not change whether mice caught crickets, but it changed how they caught them. This suggests that ZIm contributes to the structure of pursuit, not simply its initiation or outcome, actively preserving rather than improving behavioral performance under difficult conditions by shaping strategies animals use to achieve their goals^24^. Future research should examine whether this persistence function extends to other behaviors that demand sustained engagement when sensory evidence is incomplete, and which ZIm output pathways mediate these effects. By examining how behavioral distributions shift rather than only whether performance changes, we can identify circuit contributions that summary metrics would miss. Uncovering these functions requires resolving the fine structure of behavior itself, where information about the strategies that sustain success reside in distributions of the mouse-cricket interaction that averages and summary statistics cannot capture.

## Supplemental Figures

**Figure S1:**
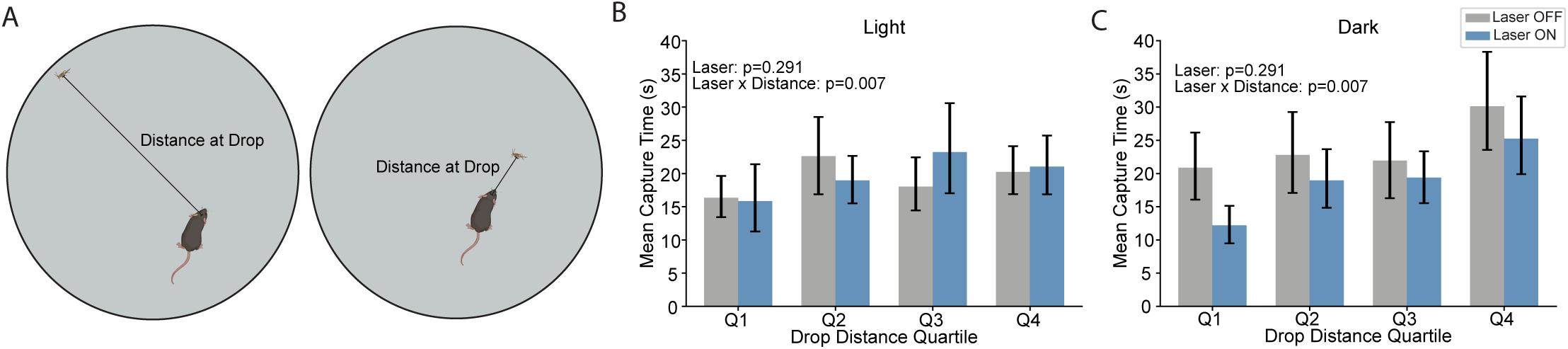
Effect of ZIm optogenetic activation depends on initial distance between the mouse and cricket. A) Example schematic of the variability of distances between the mouse and cricket at the drop in our large 60 cm diameter arena. Previous experiments^19^ were performed in a 29 cm x 15cm arena. B-C) Partitioning data based on drop distance quartiles reveals a significant interaction between laser and drop distance in the shortest drop quartile (n=7, p=0.007, Quantile Regression).

**Figure S2:**
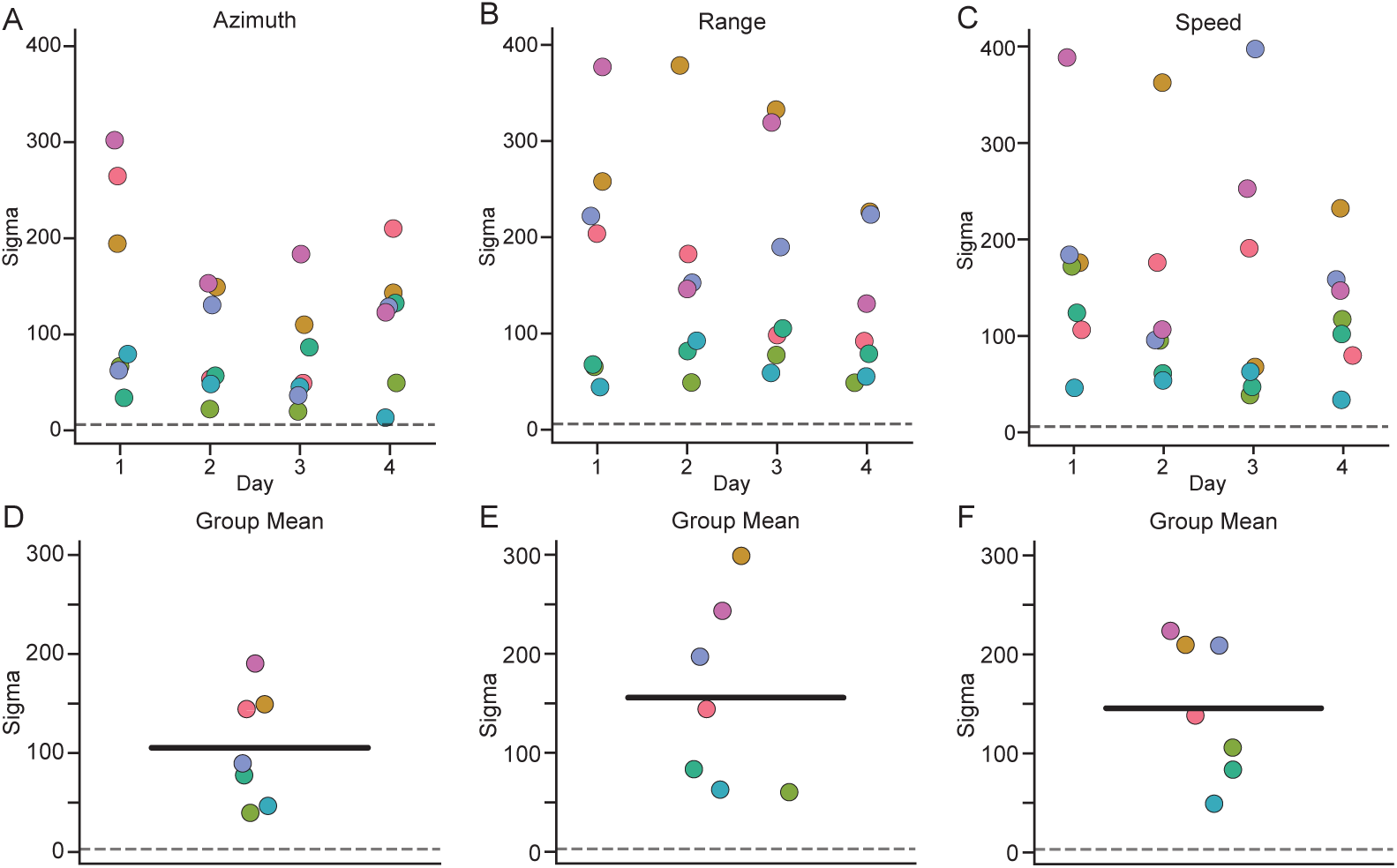
Mean sigma values for geometric distributions during baseline were computed within mouse across days. A) Individual mouse sigma values across days for azimuth. B) Individual mouse sigma values across days for range. C) Individual mouse sigma values across days for speed. D) Population mean sigma values for azimuth. E) Population mean sigma values for range. F) Population mean sigma values for speed. Dashed line indicates sigma=6, corresponding to p=0.001.

**Figure S3:**
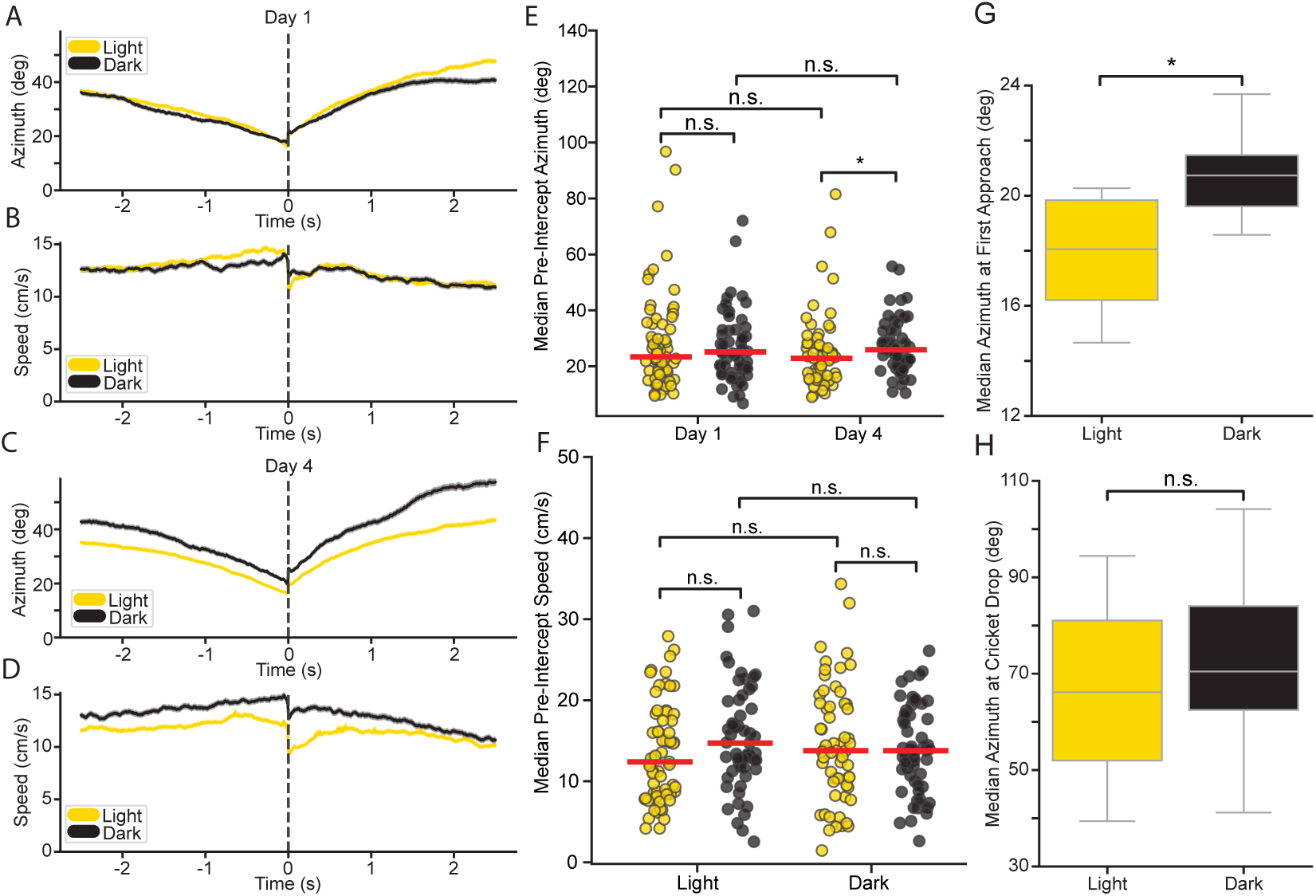
During baseline prey capture, mice develop different approach strategies in the light and dark. A) On day 1 of baseline, pre-intercept azimuth is the same in the light and dark. B) On day 1 of baseline, pre-intercept speed is the same in light and dark. C) By day 4 of baseline, pre-contact azimuth is lower in the light than in the dark. D) By day 4 of baseline, pre-intercept speed is lower in the light than in the dark. E) Median pre-intercept azimuth does not differ between light and dark on day 1 of baseline but does on day 4 (n=7; day 1: p>0.05, day 4:p=0.0465, Wilcoxon Rank Sum). F) Median pre-intercept speed does not differ between light and dark on day 1 or day 4 of baseline (n=7; all p>0.05, Wilcoxon Rank Sum). G) Median azimuth at the start of the first approach is significantly higher in the dark than in the light during baseline (n=7; p=0.016, Wilcoxon Rank Sum). H) Median azimuth at the cricket drop does not differ between light and dark during baseline (n=7; p>0.05, Wilcoxon Rank Sum.

**Figure S4:**
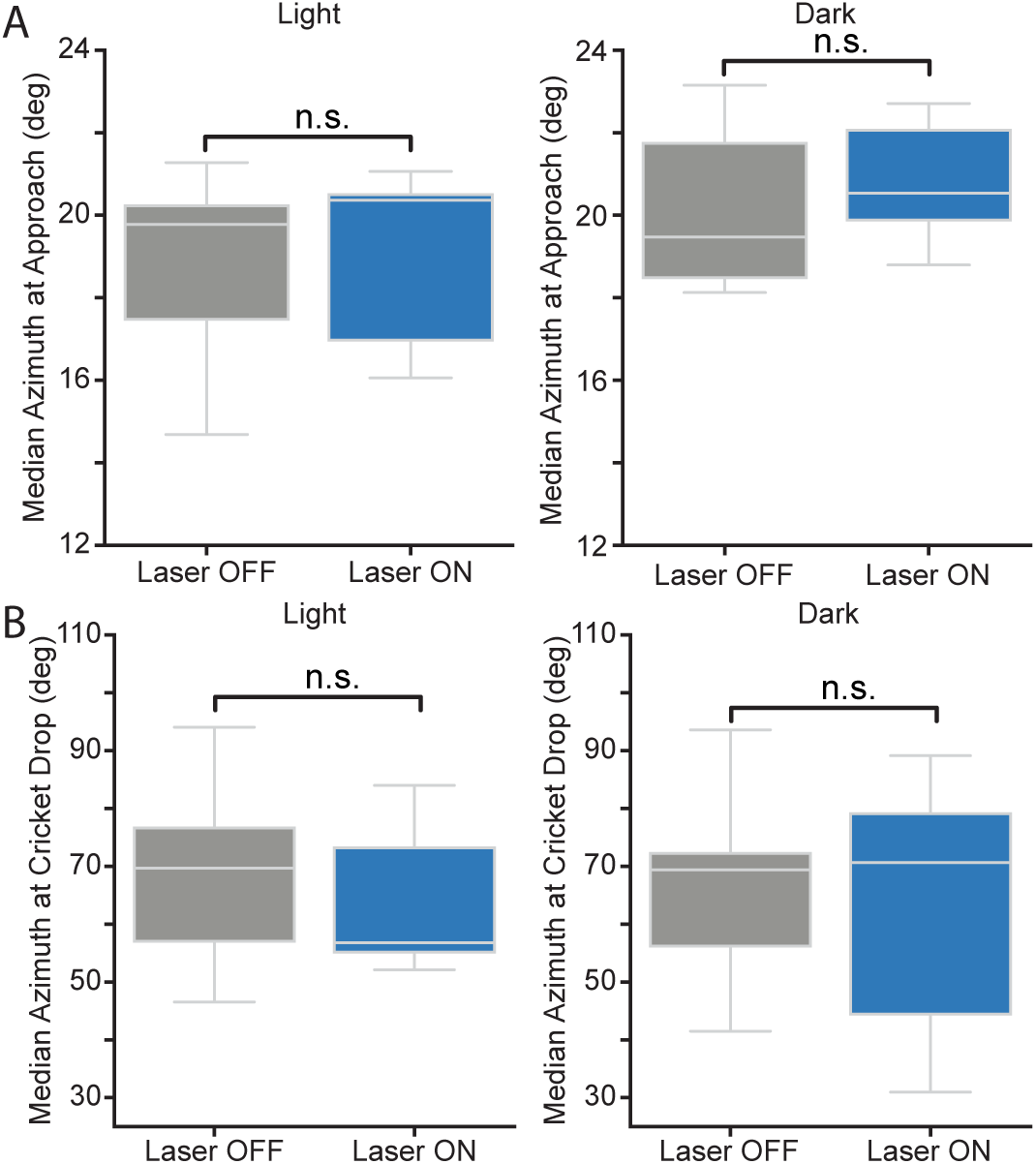
Median azimuth at the start of approach and cricket drop are not affected by optogenetic activation of ZIm. A) Median azimuth at the start of the first approach does not differ between laser on and off in the light or dark (n=7; p>0.05, Wilcoxon Rank Sum). B) Median azimuth at the cricket drop does not differ between laser on and off in the light or dark (n=7; p>0.05, Wilcoxon Rank Sum).

**Figure S5:**
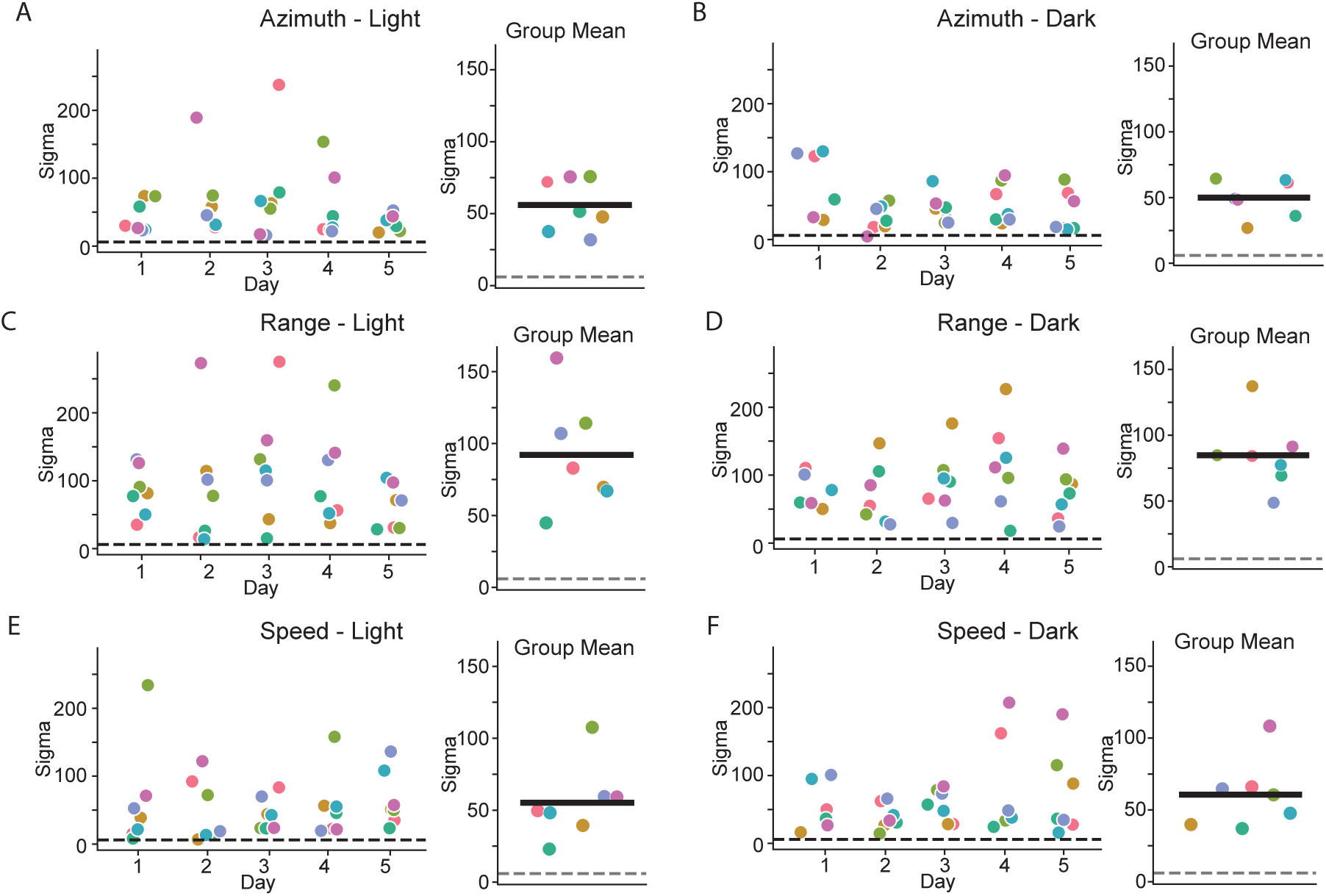
Mean sigma values for geometric distributions were computed within mouse across days. A-B) Individual mouse sigma values across days and population mean sigma for azimuth in the light and dark. C-D) Individual mouse sigma values across days and population mean sigma for range in the light and dark. E-F) Individual mouse sigma values across days and population mean sigma for mouse speed in the light and dark. Dashed line indicates sigma=6, corresponding to p=0.001.

**Figure S6:**
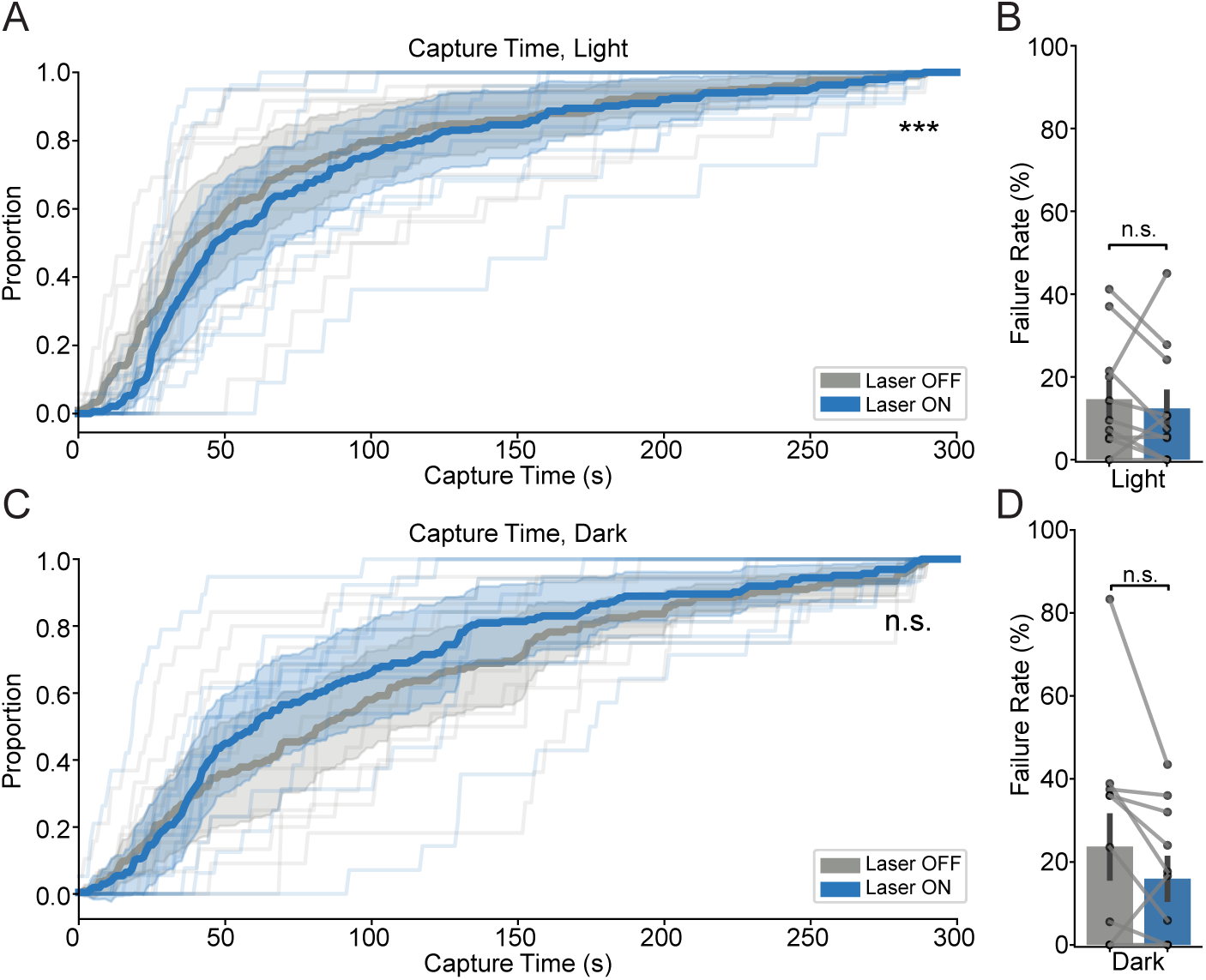
ZIm activation does not reduce capture times and failure rates in experienced viral mice. A) Capture times in the light are significantly longer with ZIm activation in the light (n=11; p<0.001, K-S test). B) Failure rate does not change with ZIm activation in the light (n=11; p>0.05, mixedLM). C) Capture times do not change with ZIm activation in the dark (n=11; p>0.05, K-S test). D) Failure rate does not change with ZIm activation in the dark (n=11; p>0.05, mixedLM).

**Figure S7:**
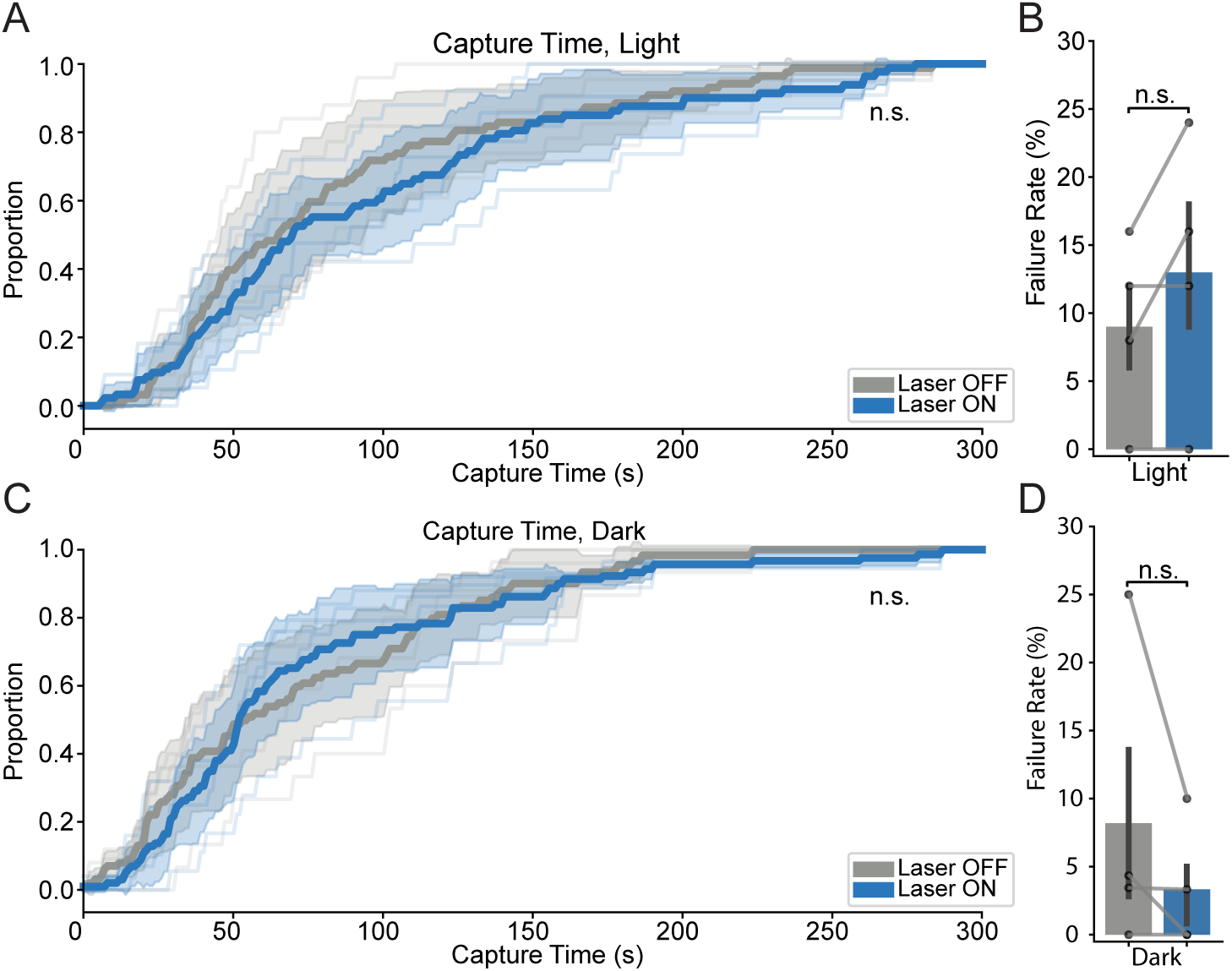
ZIm does not reduce capture times and failure rates in naïve Vgat-ChR2 mice. A) Capture times do not change with ZIm activation in the light (n=4; p>0.05, K-S test). B) Failure rate does not change with ZIm activation in the light (n=4; p>0.05, mixedLM). C) Capture times do not change with ZIm activation in the dark (n=11; p>0.05, K-S test). D) Failure rate does not change with ZIm activation in the dark (n=4; p>0.05, mixedLM).

**Figure S8:**
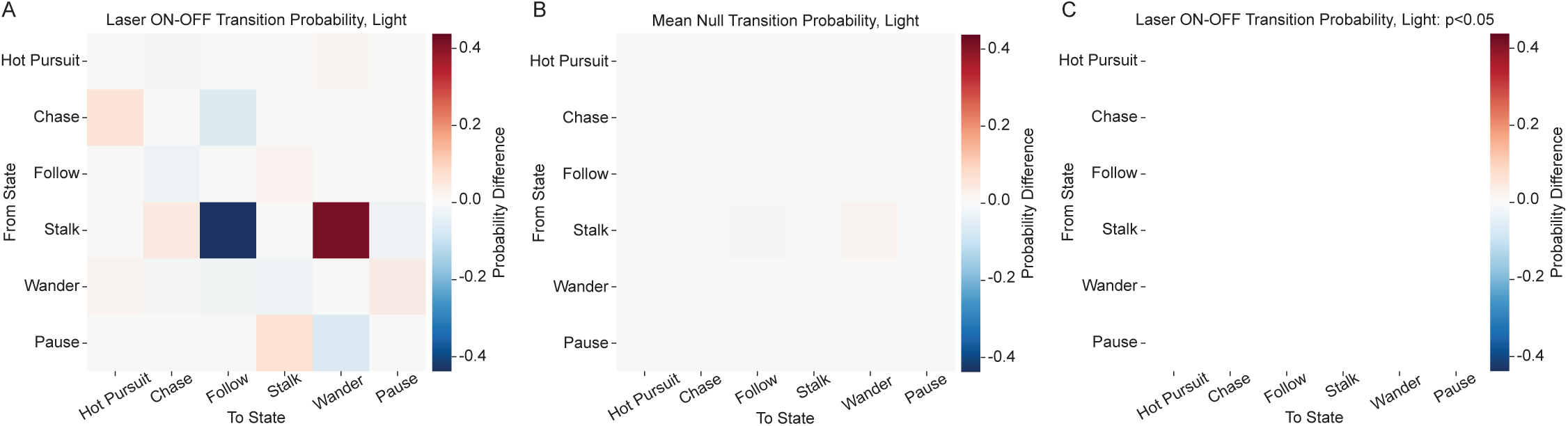
ZIm activation does not change transition structure in the light. A) Observed difference TPM between laser on and off in the light. B) Mean null partial TPM in the light generated from 10,000 permutations. C) Observed difference TPM with only significant transitions shown (n=7; all p>0.05, Permutation Test, FDR correction).

## Supplemental Tables

**Table S1:** Injection and implant coordinates.

| Brain region | AP (mm) | DV (mm) | ML (mm) | Angle (°) | Application |
|---|---|---|---|---|---|
| ZIm | -1.80 | -4.40 | +/- 1.00 | 0 (L/R) | 200 nL Virus injection |
| ZIm | -1.80 | -3.65 | +/- 1.50 | 0 (L/R) | Optic fiber implant |

**Table S2:** Permutation Test Results, Dark-Light Transitions.

| From State | To State | Observed Difference | Null Mean | p_value | FDR q_value |
|---|---|---|---|---|---|
| Follow | Stalk | 0.190715 | 0.006119 | 0.000100 | 0.000900 |
| Follow | Chase | -0.189407 | -0.006283 | 0.000100 | 0.000900 |
| Chase | Follow | 0.205580 | 0.012319 | 0.000100 | 0.000900 |
| Chase | Hot Pursuit | -0.225579 | -0.013093 | 0.000100 | 0.000900 |
| Stalk | Wander | 0.327626 | 0.013455 | 0.003200 | 0.023038 |
| Wander | Stalk | 0.163787 | 0.008144 | 0.006399 | 0.038396 |
| Stalk | Follow | -0.270548 | -0.000644 | 0.010399 | 0.053480 |
| Wander | Pause | -0.170100 | -0.006967 | 0.018098 | 0.081442 |
| Chase | Stalk | 0.019999 | 0.000774 | 0.033197 | 0.132787 |
| Pause | Wander | -0.068632 | 0.003423 | 0.043996 | 0.158384 |
| Pause | Stalk | 0.056132 | -0.002647 | 0.094791 | 0.310224 |
| Pause | Follow | 0.012500 | -0.000776 | 0.163484 | 0.490451 |
| Wander | Wander | 0.000000 | 0.000000 | 1.000000 | 1.000000 |
| Pause | Hot Pursuit | 0.000000 | 0.000000 | 1.000000 | 1.000000 |
| Stalk | Stalk | 0.000000 | 0.000000 | 1.000000 | 1.000000 |
| Wander | Follow | 0.004208 | -0.001576 | 0.813919 | 1.000000 |
| Wander | Chase | 0.002104 | 0.000399 | 0.757324 | 1.000000 |
| Pause | Chase | 0.000000 | 0.000000 | 1.000000 | 1.000000 |
| Wander | Hot Pursuit | 0.000000 | 0.000000 | 1.000000 | 1.000000 |
| Stalk | Pause | -0.014840 | -0.008597 | 0.809319 | 1.000000 |
| Hot Pursuit | Hot Pursuit | 0.000000 | 0.000000 | 1.000000 | 1.000000 |
| Follow | Pause | 0.000000 | 0.000000 | 1.000000 | 1.000000 |
| Stalk | Hot Pursuit | 0.000000 | 0.000000 | 1.000000 | 1.000000 |
| Follow | Wander | -0.001308 | 0.000165 | 0.704630 | 1.000000 |
| Follow | Follow | 0.000000 | 0.000000 | 1.000000 | 1.000000 |
| Follow | Hot Pursuit | 0.000000 | 0.000000 | 1.000000 | 1.000000 |
| Chase | Pause | 0.000000 | 0.000000 | 1.000000 | 1.000000 |
| Chase | Wander | 0.000000 | 0.000000 | 1.000000 | 1.000000 |
| Chase | Chase | 0.000000 | 0.000000 | 1.000000 | 1.000000 |
| Hot Pursuit | Pause | 0.000000 | 0.000000 | 1.000000 | 1.000000 |
| Hot Pursuit | Wander | 0.000000 | 0.000000 | 1.000000 | 1.000000 |
| Hot Pursuit | Stalk | 0.000000 | 0.000000 | 1.000000 | 1.000000 |
| Hot Pursuit | Follow | 0.000000 | 0.000000 | 1.000000 | 1.000000 |
| Hot Pursuit | Chase | 0.000000 | 0.000000 | 1.000000 | 1.000000 |
| Stalk | Chase | -0.042237 | -0.004214 | 0.409659 | 1.000000 |
| Pause | Pause | 0.000000 | 0.000000 | 1.000000 | 1.000000 |

**Table S3:** Permutation Test Results, Laser ON-OFF Light.

| From State | To State | Observed Difference | Null Mean | p_value | FDR q_value |
|---|---|---|---|---|---|
| Stalk | Follow | -0.437500 | -0.009600 | 0.011799 | 0.277172 |
| Stalk | Wander | 0.416667 | 0.012332 | 0.015398 | 0.277172 |
| Pause | Stalk | 0.065639 | -0.001098 | 0.053795 | 0.484152 |
| Pause | Wander | -0.065639 | 0.001098 | 0.053795 | 0.484152 |
| Hot Pursuit | Chase | -0.008696 | 0.000435 | 0.180282 | 0.942306 |
| Hot Pursuit | Wander | 0.008696 | -0.000435 | 0.180282 | 0.942306 |
| Chase | Hot Pursuit | 0.058016 | 0.002739 | 0.235576 | 0.942306 |
| Chase | Follow | -0.058542 | -0.002528 | 0.224678 | 0.942306 |
| Wander | Hot Pursuit | 0.012195 | -0.000382 | 0.220878 | 0.942306 |
| Stalk | Pause | -0.020833 | -0.006117 | 0.853915 | 1.000000 |
| Wander | Chase | -0.009524 | -0.000950 | 0.739726 | 1.000000 |
| Wander | Follow | -0.019048 | -0.001104 | 0.352465 | 1.000000 |
| Hot Pursuit | Hot Pursuit | 0.000000 | 0.000000 | 1.000000 | 1.000000 |
| Wander | Wander | 0.000000 | 0.000000 | 1.000000 | 1.000000 |
| Stalk | Stalk | 0.000000 | 0.000000 | 1.000000 | 1.000000 |
| Pause | Hot Pursuit | 0.000000 | 0.000000 | 1.000000 | 1.000000 |
| Pause | Chase | 0.000000 | 0.000000 | 1.000000 | 1.000000 |
| Pause | Follow | 0.000000 | 0.000000 | 1.000000 | 1.000000 |
| Wander | Stalk | -0.020557 | 0.000466 | 0.521148 | 1.000000 |
| Wander | Pause | 0.036934 | 0.001970 | 0.378862 | 1.000000 |
| Follow | Pause | 0.000000 | 0.000000 | 1.000000 | 1.000000 |
| Stalk | Hot Pursuit | 0.000000 | 0.000000 | 1.000000 | 1.000000 |
| Follow | Wander | 0.002462 | -0.000097 | 0.674933 | 1.000000 |
| Follow | Stalk | 0.020374 | -0.000274 | 0.525847 | 1.000000 |
| Follow | Follow | 0.000000 | 0.000000 | 1.000000 | 1.000000 |
| Follow | Chase | -0.022835 | 0.000371 | 0.493351 | 1.000000 |
| Follow | Hot Pursuit | 0.000000 | 0.000000 | 1.000000 | 1.000000 |
| Chase | Pause | 0.000000 | 0.000000 | 1.000000 | 1.000000 |
| Chase | Wander | 0.000000 | 0.000000 | 1.000000 | 1.000000 |
| Chase | Stalk | 0.000526 | -0.000210 | 0.767723 | 1.000000 |
| Chase | Chase | 0.000000 | 0.000000 | 1.000000 | 1.000000 |
| Hot Pursuit | Pause | 0.000000 | 0.000000 | 1.000000 | 1.000000 |
| Hot Pursuit | Stalk | 0.000000 | 0.000000 | 1.000000 | 1.000000 |
| Hot Pursuit | Follow | 0.000000 | 0.000000 | 1.000000 | 1.000000 |
| Stalk | Chase | 0.041667 | 0.003385 | 0.765523 | 1.000000 |
| Pause | Pause | 0.000000 | 0.000000 | 1.000000 | 1.000000 |

**Table S4:** Permutation Test Results, Laser ON-OFF Dark.

| From State | To State | Observed Difference | Null Mean | p_value | FDR q_value |
|---|---|---|---|---|---|
| Chase | Hot Pursuit | 0.226097 | 0.004704 | 0.000100 | 0.001800 |
| Chase | Follow | -0.214720 | -0.005161 | 0.000100 | 0.001800 |
| Stalk | Wander | -0.315721 | -0.003045 | 0.003600 | 0.043196 |
| Stalk | Follow | 0.234834 | 0.000611 | 0.023998 | 0.215978 |
| Follow | Stalk | -0.120169 | 0.001392 | 0.034897 | 0.251255 |
| Follow | Chase | 0.117580 | -0.001516 | 0.052795 | 0.316768 |
| Stalk | Chase | 0.101761 | 0.001974 | 0.063994 | 0.329110 |
| Pause | Wander | 0.087500 | 0.000389 | 0.109789 | 0.494051 |
| Pause | Stalk | -0.075000 | -0.001335 | 0.157484 | 0.629937 |
| Wander | Pause | 0.125379 | -0.002063 | 0.226477 | 0.815318 |
| Stalk | Stalk | 0.000000 | 0.000000 | 1.000000 | 1.000000 |
| Stalk | Pause | -0.020874 | 0.000459 | 0.682132 | 1.000000 |
| Hot Pursuit | Hot Pursuit | 0.000000 | 0.000000 | 1.000000 | 1.000000 |
| Wander | Chase | -0.011628 | -0.000438 | 0.999700 | 1.000000 |
| Wander | Stalk | -0.090495 | 0.002272 | 0.320168 | 1.000000 |
| Wander | Wander | 0.000000 | 0.000000 | 1.000000 | 1.000000 |
| Pause | Hot Pursuit | 0.000000 | 0.000000 | 1.000000 | 1.000000 |
| Pause | Chase | 0.000000 | 0.000000 | 1.000000 | 1.000000 |
| Pause | Follow | -0.012500 | 0.000946 | 0.998700 | 1.000000 |
| Wander | Hot Pursuit | 0.000000 | 0.000000 | 1.000000 | 1.000000 |
| Wander | Follow | -0.023256 | 0.000229 | 0.497850 | 1.000000 |
| Follow | Pause | 0.000000 | 0.000000 | 1.000000 | 1.000000 |
| Follow | Wander | 0.002589 | 0.000125 | 0.538946 | 1.000000 |
| Follow | Follow | 0.000000 | 0.000000 | 1.000000 | 1.000000 |
| Follow | Hot Pursuit | 0.000000 | 0.000000 | 1.000000 | 1.000000 |
| Chase | Pause | 0.000000 | 0.000000 | 1.000000 | 1.000000 |
| Chase | Wander | 0.000000 | 0.000000 | 1.000000 | 1.000000 |
| Chase | Stalk | -0.011377 | 0.000457 | 0.353365 | 1.000000 |
| Chase | Chase | 0.000000 | 0.000000 | 1.000000 | 1.000000 |
| Hot Pursuit | Pause | 0.000000 | 0.000000 | 1.000000 | 1.000000 |
| Hot Pursuit | Wander | 0.000000 | 0.000000 | 1.000000 | 1.000000 |
| Hot Pursuit | Stalk | 0.000000 | 0.000000 | 1.000000 | 1.000000 |
| Hot Pursuit | Follow | 0.000000 | 0.000000 | 1.000000 | 1.000000 |
| Hot Pursuit | Chase | 0.000000 | 0.000000 | 1.000000 | 1.000000 |
| Stalk | Hot Pursuit | 0.000000 | 0.000000 | 1.000000 | 1.000000 |
| Pause | Pause | 0.000000 | 0.000000 | 1.000000 | 1.000000 |

